# Antigen Presentation-Independent Reciprocal Immune Modulation by *HLA-DRB1* Allelic Epitopes that Associate with Autoimmune Disease Risk or Protection

**DOI:** 10.1101/2020.08.25.265348

**Authors:** Vincent van Drongelen, Bruna Miglioranza Scavuzzi, Sarah Veloso Nogueira, Frederick W. Miller, Amr H. Sawalha, Joseph Holoshitz

## Abstract

Statistical associations between particular human leukocyte antigen (HLA) alleles and susceptibility to - or protection from - autoimmune diseases have been long observed. Allele-specific antigen presentation (AP) has been widely proposed as a culprit mechanism; however, direct evidence to substantiate that hypothesis is scant. Here we demonstrate AP-independent differential macrophage activation by *HLA-DRB1* alleles known to associate with autoimmune disease risk or protection with resultant polarization of pro-inflammatory (“M1”) versus anti-inflammatory (“M2”) macrophages, respectively. RNA-sequencing analyses of *in vitro*-polarized macrophages in the presence of AP-incompetent short synthetic peptides corresponding to the third allelic hypervariable regions coded by those two *HLA-DRB1* alleles showed reciprocal activation of pro- versus anti-inflammatory transcriptomes, with implication of corresponding gene ontologies and upstream regulators. These results identify a previously unrecognized mechanism of differential immune modulation by short *HLA-DRB1*-coded allelic epitopes independent of AP, and could shed new light on the mechanistic basis of HLA-disease association.

## Introduction

It has been long observed that certain human leukocyte antigen (HLA) alleles confer higher risk for autoimmune diseases, while other alleles provide protective effects^1^. The molecular mechanisms underlying these associations are incompletely understood. Modeled on the major histocompatibility complex (MHC)-restricted antigen presentation (AP) paradigm^2, 3^, it has been hypothesized that HLA-associated diseases are triggered by presentation of self-antigens - or foreign antigens that cross-react with self by HLA-coded molecules^4, 5, 6, 7^. It is worth noting, however, that decades after the AP-based hypotheses have been invoked, the identity of the putative target antigens remains unknown in most HLA-associated conditions. Furthermore, as previously discussed^8, 9^, the AP-based hypotheses are difficult to reconcile with the promiscuous associations of particular HLA alleles with many unrelated diseases.

For example, although the majority of individuals with rheumatoid arthritis (RA) carry *HLA-DRB1* alleles that code for a susceptibility (or ‘shared’) epitope (SE) motif QKRAA, QRRAA, or RRRAA in position 70-74 of the DRβ chain^10^, the association is not disease-specific. SE-coding *HLA-DRB1* alleles have also been found to associate with type 1 diabetes, autoimmune hepatitis, polymyalgia rheumatica, and temporal arteritis^11, 12, 13, 14^, among other conditions. Additionally, the SE has been found to be a risk factor for erosive bone damage in nosologically-distinct conditions, such as periodontal disease, systemic lupus erythematosus and psoriatic arthritis^15, 16, 17, 18, 19^.

Inversely, *HLA-DRB1* alleles that encode a protective epitope (PE) motif 70-DERAA-74 (at the exact same region of the DRβ chain as the SE) significantly reduce RA risk^20, 21, 22, 23^, as well as many other autoimmune conditions. Examples include systemic lupus erythematosus, mixed connective tissue disease, progressive systemic sclerosis, antineutrophil cytoplasmic antibodies (ANCA)-positive vasculitis, narcolepsy and myasthenia gravis^24, 25, 26, 27, 28, 29^. Thus, both the SE and PE modulate disease risk in a wide range of conditions that do not share a common putative target antigen, pathogenesis, or target tissues. Such infidelity is difficult to reconcile with basic tenets of the MHC-restricted AP paradigm.

To determine whether the SE and PE effects may be AP-independent, here we compared their respective impacts on macrophage activation, given the pivotal role that these cells play in autoimmunity^30, 31, 32, 33, 34^. The findings demonstrate that primary bone marrow macrophages from naïve transgenic mice that constitutively express on their cell surface SE-positive HLA-DR molecules are polarized preferentially toward pro-inflammatory (“M1”) macrophage, while macrophages from transgenic mice that express PE-positive HLA-DR molecules show enhanced anti-inflammatory (“M2”) polarization. To exclude the possibility of AP of self- or tissue culture-derived antigens by the transgenic HLA-DR molecules, and to map the functional epitopes, we studied AP-incompetent soluble synthetic 15-mer peptides corresponding to the third allelic hypervariable regions (TAHRs) coded by these two *HLA-DRB1* alleles. Using an RNA sequencing (RNA-seq) approach, we demonstrate here that consistent with the above findings, exposure of macrophages to SE- or PE-expressing peptides activates distinct transcriptomes. A SE-expressing peptide activates genes, biological processes and upstream regulators known to mediate pro-M1, pro-inflammation or pro-autoimmune disease effects, whereas a PE-expressing peptide facilitates activation of genes, biological processes and upstream regulators that are known to mediate pro-M2, anti-inflammatory, or anti-autoimmune disease effects. Thus, gene products coded by *HLA-DRB1* alleles that are known to associate with autoimmune disease susceptibility versus protection activate, respectively, pro-inflammatory or anti-inflammatory pathways in an AP-independent fashion.

## Results

### HLA-DRB1 allele-specific differential macrophage polarization in transgenic mice

To determine whether autoimmune disease risk or protective *HLA-DRB1* alleles have distinct effects on macrophage polarization we first studied *ex vivo* primary bone marrow-derived macrophages (BMDMs) isolated from transgenic mice^35, 36^ that express human HLA-DRβ chains coded by the *DRB1*04:02* allele with a PE (70-DERAA-74) sequence in the TAHR (“PE Tg”), versus BMDMs from mice of the same genetic background expressing a SE (70-QKRAA-74) sequence, encoded by the RA-susceptibility allele *DRB1*04:01* (“SE Tg”). The TAHRs of the two transgenic mouse lines differ by only 3 amino acid residues. As shown in Fig. 1, under M1-polarizing culture conditions, SE Tg-derived BMDMs expressed higher levels of the M1 gene markers *Cxcl10, Nos2, Il-12p40, Il-1b, Tnfa, Il-6* and *Ccl2* compared to PE Tg-derived BMDMs (Fig. 1A). Additionally, SE Tg-derived BMDMs produced Il-6, Tnfα, and Il-12p70 proteins at significantly higher levels than PE Tg-derived BMDMs (Fig. 1B). SE Tg BMDMs also showed higher baseline levels of nitric oxide (NO) production with a significant increase under M1 polarizing culture conditions, whereas PE Tg-derived BMDMs did not produce any NO in either culture conditions (Fig. 1C). A mirror-image pattern was found under M2-polarizing culture conditions: PE Tg BMDMs demonstrated higher expression of the M2 gene markers *Arg1, Ym1* and *Ccl17* relative to SE Tg BMDMs (Fig. 1D). Consistent with the above findings, and with the role of arginase in M2 polarization^37^, a significantly increased arginase activity under M2 polarizing conditions was found in BMDMs from PE Tg, compared to SE Tg (Fig. 1E).

**Fig. 1.**
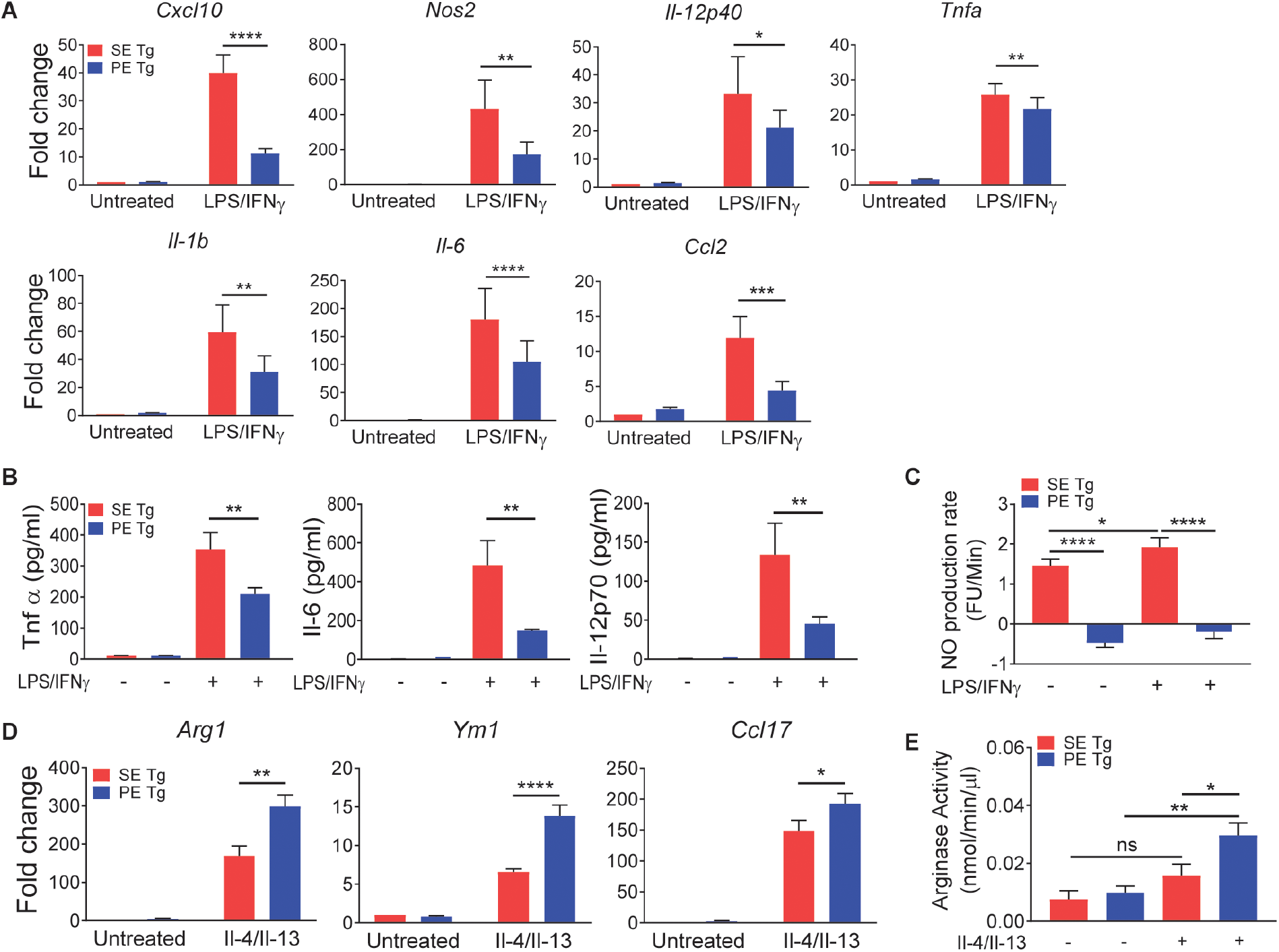
Differential in vitro macrophage polarization in SE Tg and PE Tg BMDMs. For M1 polarization, BMDMs were treated with LPS (1 ng/ml) + IFNγ (20 ng/ml) for 24 hrs. M2 polarization was induced with Il-4 (10 ng/ml) + Il-13 (10 ng/ml) for 24 hrs. (**A**) qPCR for M1-associated genes. Data represent mean + SEM of 3-5 independent experiments. (**B**) ELISA for M1-associated cytokines. Data represent mean + SEM of 3 independent experiments. (**C**) NO production by BMDMs under M1 polarization conditions. Data represent mean + SEM of 8 replicates from 2 independent experiments. (**D**) qPCR for M2-associated genes. Data represent mean + SEM of 3-4 independent experiments. (**E**) Arginase activity in BMDMs under M2 polarization conditions. Data represent mean + SEM of 3 independent experiments. 2-way ANOVA, *P<0.05, **P<0.01, ***P<0.001, ****P<0.0001

*HLA-DRB1* allele-based differential macrophage polarization was found *in vivo* as well. As shown in Fig. 2A, intra peritoneal (i.p.) administration of an M1-polarizing agent (LPS) significantly increased gene expression of M1 markers *Il12p40, Il12p35, Il23p19, Il-1b, Tnfa, Ccl2* and *Il-6* in peritoneal macrophages from SE Tg, compared to PE Tg mice. Additionally, SE Tg mice produced higher serum levels of various pro-inflammatory, M1-associated cytokines, including Il-12, TNFα, GM-CSF and IFNγ (Fig. 2B).

**Fig. 2.**
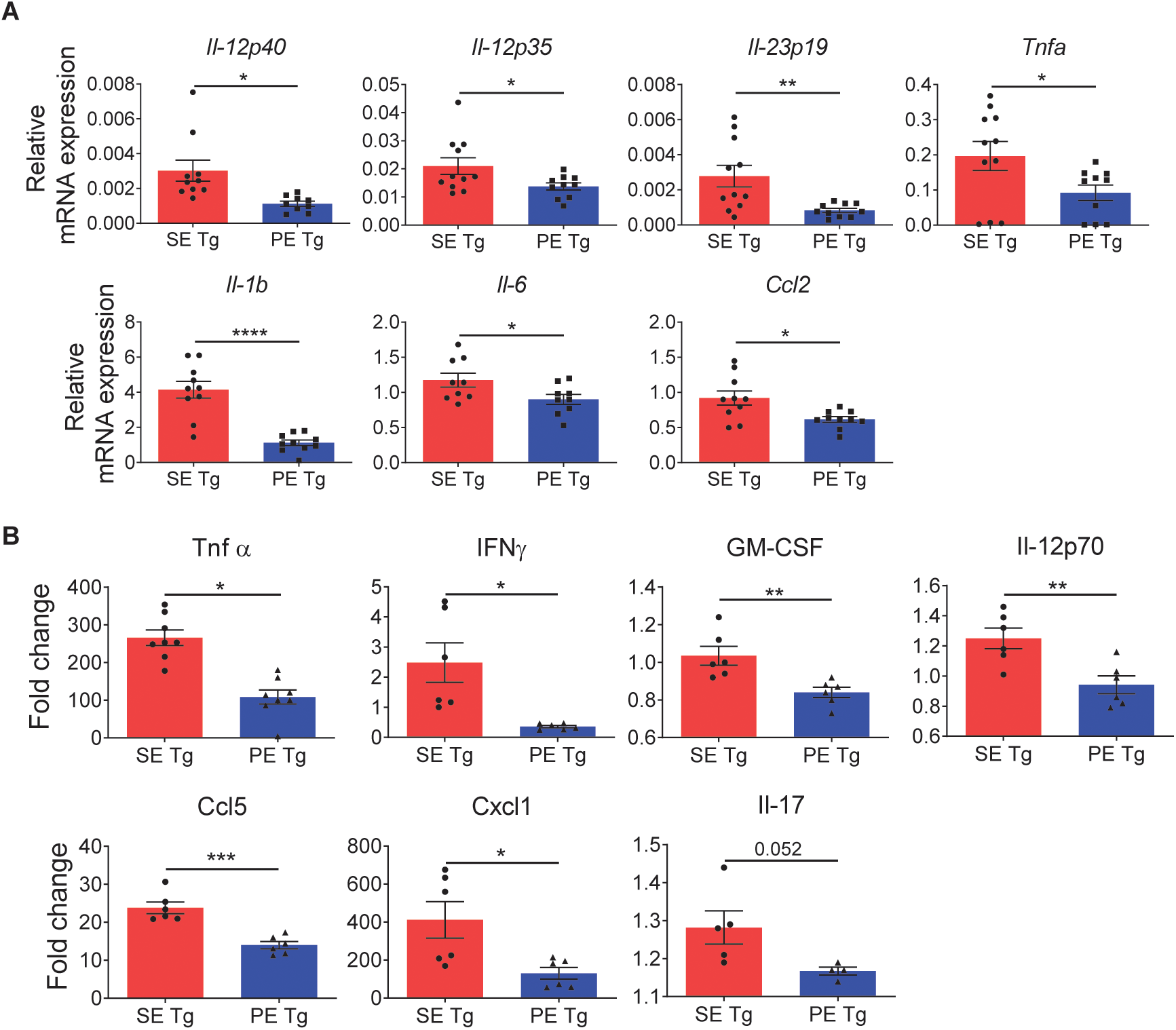
Differential in vivo macrophage polarization in SE Tg and PE Tg mice. To induce M1 polarization *in vivo*, mice were injected i.p. with LPS (500 μg/kg). Peritoneal macrophages and serum were collected after 4 hours. (**A**) qPCR for mRNA expression of M1 marker genes in peritoneal macrophages. n=8-11 (**B**) ELISA for serum cytokine levels. n=5-8. Results are compiled data from 3 experiments. Mean ± SEM, unpaired *t*-test with welch correction, *P<0.05, **P<0.01, ***P<0.001, ****P<0.0001

### Signal transduction pathways

To better understand the mechanisms involved in *HLA-DRB1* allele-based macrophage polarization predilections, we sought to determine whether signaling events are involved. To this end we focused on the Akt axis, known to play a pivotal role in M1 versus M2 macrophage polarization^38, 39^. As Fig. 3 shows, under M1 polarizing conditions, BMDMs from PE Tg mice displayed significantly increased Akt phosphorylation compared to BMDMs from SE Tg mice (Fig. 3A). Ly294002-mediated inhibition of Pi3K, an upstream regulator of Akt, allowed significant rescue of the expression of M1 gene markers *Il-1b* and *Il-6* (Fig. 3B), as well as Il-6 and Tnfα cytokine levels (Fig. 3C). To better characterize the upstream signaling events that impact Akt activation, we measured phosphorylation of two Pi3K regulating phosphatases: PTEN and SHIP1 under M1 polarizing conditions. PTEN phosphorylation levels were not different between SE Tg and PE Tg BMDMs (Fig. S1A). However, SHIP1, showed significantly lower phosphorylation levels in PE Tg BMDMs compared to SE Tg (Fig. S1B).

**Fig. 3.**
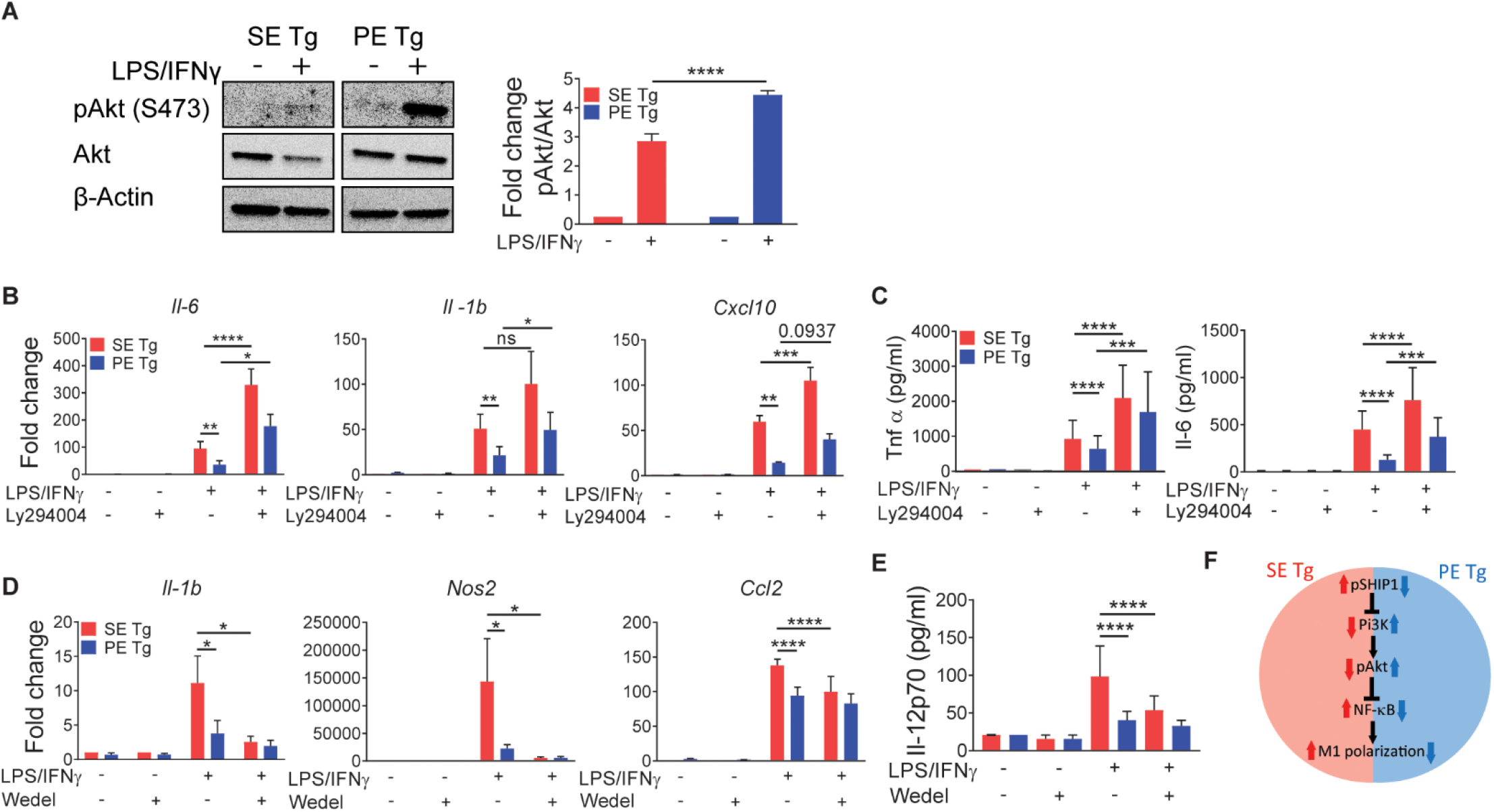
Involvement of signaling pathways in SE Tg and PE Tg BMDMs under M1 polarization conditions. M1 polarization in BMDMs was performed using the conditions described in Fig. 1. (**A**) Immunoblot for pAKt (Ser473) and Akt 15 min. after exposure of cells to M1 polarization conditions. Quantification data represent mean + SEM of 3 independent experiments. (**B** to **E**) BMDMs were pre-treated with Ly294002 (5 μM) (B,C), or wedelolactone (10 μM) (D,E) for 1 h., followed by incubation in M1 polarization conditions as in (A). (**B** and **D**) qPCR-based determination of M1 gene marker expression levels. (**C** and **E**) ELISA for Tnfa or Il-6 (C), or IL12p70 (E). (**F**) A proposed model of signaling pathway involvement under M1 polarizing conditions. Data represent mean and SEM of 3-5 independent experiments. Statistics: Within group comparisons, paired *t*-test; between groups comparison, 2-way ANOVA, *P<0.05, **P<0.01, ***P<0.001, ****P<0.0001.

Another important signaling mechanism in M1 polarization is NF-κB^40^, mapped downstream of Akt^41^. We therefore asked whether NF-κB plays a role in *HLA-DRB1*-associated macrophage polarization. Under M1 polarizing conditions, inhibition of NF-κB decreased expression of M1 gene markers *Il-1b, Nos2* and *Ccl2*, as well as the M1 cytokine Il-12p70 in SE Tg BMDMs, but not in PE Tg BMDMs (Fig. 3D and E). Thus, taken together, we propose that the diminished M1 polarizability of PE Tg BMDMs is secondary to increased Akt activation, previously shown to inhibit NF-κB signaling^38^. We further propose that increased SHIP1 activity in SE Tg BMDMs results in reduced Akt activation, leading to enhanced NF-κB activity, which, in turn leads to increased M1 polarization, consistent with previous studies^38^. A proposed model of the signaling pathways involved in *HLA-DRB1* allele-specific M1 macrophage polarization is shown in Fig. 3F.

Under M2 polarization conditions, significantly higher Akt phosphorylation was found in PE Tg BMDMs compared to SE Tg BMDMs (Fig. 4A). Modulation of the Akt signaling pathway through inhibition of Pi3K (upstream of Akt), or p70S6K (downstream of Akt), significantly suppressed expression of the M2 gene marker *Arg1* (Fig. 4B and 4C, respectively). Noteworthy, another M2 gene marker *Ym1* was not affected by either Pi3K or p70S6K inhibitors, suggesting that, different from *Arg1*, increased *Ym1* gene expression in PE Tg BMDMs is Akt-independent. The possibility that *Ym1* gene expression in PE Tg BMDMs is controlled by Stat6, as previously reported in other cells^42^, is consistent with significantly higher Stat6 phosphorylation levels under M2 polarization conditions in PE Tg, compared to SE Tg BMDMs (Fig. S2). A proposed model of the signaling pathways involved in *HLA-DRB1* allele-specific M2 macrophage polarization is shown in Fig. 4D.

**Fig. 4.**
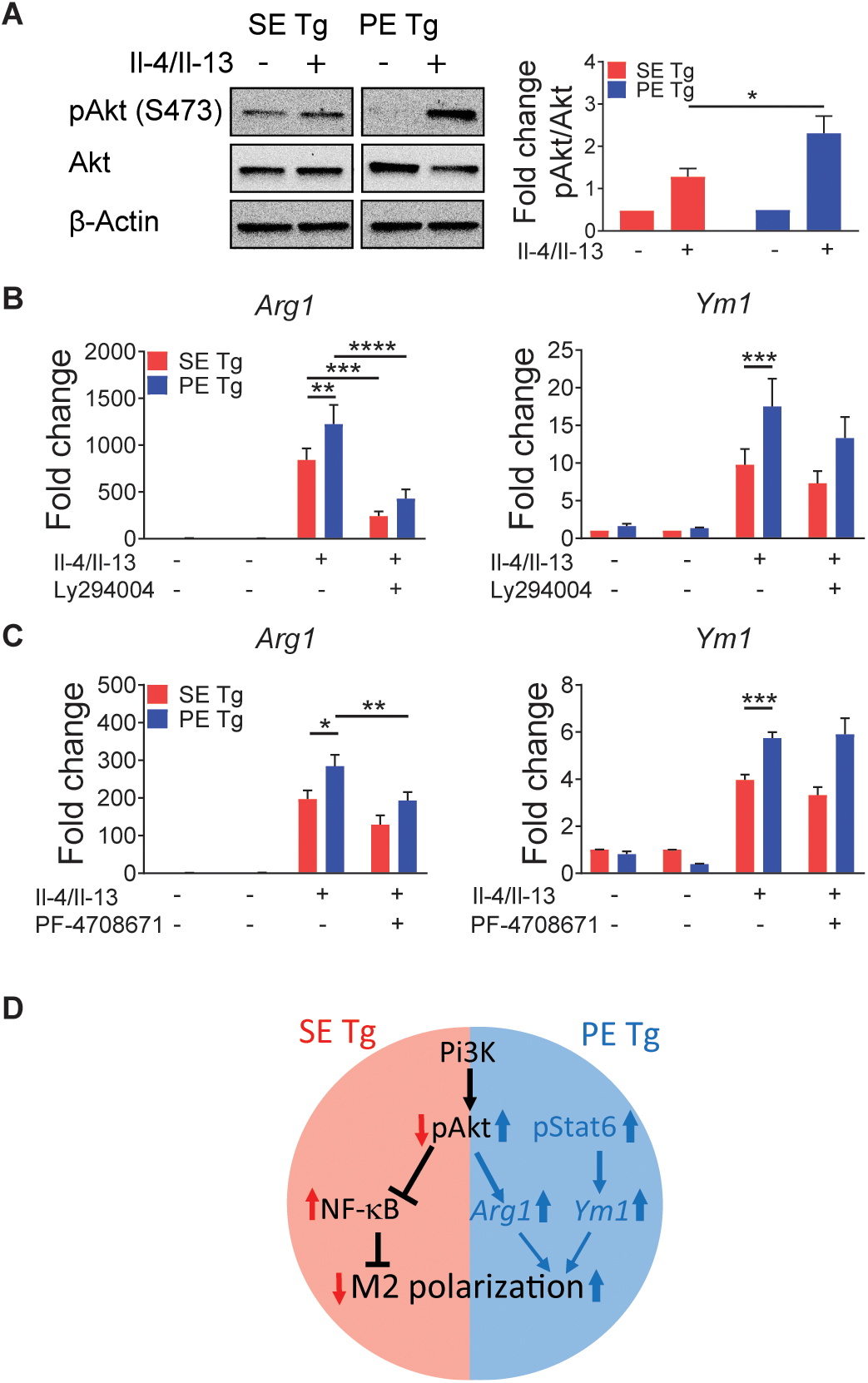
Involvement of signaling pathways in SE Tg and PE Tg BMDMs under M2 polarization conditions. M2 polarization of BMDMs was performed as in Fig. 1. (**A**) Immunoblotting for pAKt (Ser473) and Akt in BMDMs 10 min after exposure of cells to M2 polarization conditions. Quantification data represent mean + SEM of 3 independent experiments. (**B** and **C**) qPCR analysis for M2 gene markers *Arg1* and *Ym1* expression in BMDMs pre-treated with Ly294002 (5 μM) (B), or PF4708671 (10 μM) (C) for 1 hr., followed by M2 polarization for 20 hrs. Data represent mean +SEM of 3 independent experiments. Statistics: Within group comparisons, paired *t*-test; between groups comparison, 2-way ANOVA, *P<0.05, **P<0.01, ***P<0.001, ****P<0.0001. (**D**) A proposed model of signaling pathway involvement under M2 polarizing conditions.

### HLA-DRB1 allele-specific transcriptome activation

To more conclusively determine whether the differential effects of the two *HLA-DRB1* alleles may be AP-independent and to determine whether their effects could be mapped to the TAHR, we used AP-incompetent 15-mer synthetic peptides corresponding to the TAHR of the DRβ chain. To explore the feasibility of this approach, we first quantified expression levels of macrophage polarization marker genes in RAW 264.7 mouse macrophages following exposure to 15-mer peptides designated “65-79*SE” or “65-79*PE”, corresponding to amino acid residues 65-79 (TAHR) coded, respectively, by susceptibility (*HLA-DRB1*04:01*) or protective (*e.g. HLA-DRB1*04:02, DRB1*13:01*, or *DRB1*13:02*) alleles. The 65-79*SE and 65-79*PE peptides differ by 3 amino acid residues, including only 2 substitutions in the 70-74 region (QKRAA versus DERAA). Quantitative RT-PCR data (Fig. S3) confirmed that under M1-polarizing conditions, 65-79*SE, but not the 65-79*PE, activated transcription of the M1-marker genes *Nos1, Cxcl10, Ccl2* and *Il-1b* (Fig. S3A). Under M2 polarizing conditions, 65-79*PE selectively upregulated expression of the M2 marker gene *Mgl2* (Fig. S3B). Thus, consistent with the findings in transgenic mouse BMDMs, 15-mer peptides 65-79*SE and 65-79*PE, corresponding to the TAHRs coded by *DRB1* alleles that confer autoimmune disease risk or protection, respectively, recapitulated the differentially induced expression of M1 versus M2 macrophage polarization gene markers in an epitope-specific fashion.

We next used an RNA-seq approach to characterize the broader transcriptional effects of the two TAHRs in mouse RAW 264.7 macrophages stimulated with 65-79*SE or 65-79*PE. Under M1 polarizing conditions (Fig. 5) the two TAHR peptides had a distinct effect on the number of upregulated and downregulated differentially expressed genes (DEGs) (Fig. 5A and data file S1A). Among the unique upregulated genes induced by 65-79*SE in M1-polarizing conditions (Fig. 5B and C, Table S1, and data file S2A) were many RA disease marker - or risk factor - genes (e.g. *Stat3, Cd44, Traf1, Tnfaip3*), genes that encode confirmed or proposed therapeutic targets (e.g. *Jak1, Jak3, Stat3, Ccr1, Cxcr4, Mmp14*), and genes involved in autoimmune disease pathogenic mechanisms, such as NF-κB activation (e.g. *Relb, Kpna4, Vav1*), angiogenesis (*Dusp4, Pgf, Egr1, Vegfc, Vegfa, Hbegf, Hmga1*), Th17 polarization (e.g. *Dusp4, Runx1, Hbegf*), osteoclastogenesis (e.g. *Atf4, Adam8, Dcstamp, Cxcr1*), or M1 polarization (*Klf6*). Conversely, 65-79*SE downregulated many anti-inflammatory (e.g. *Akap1, Casp9, Cyp51, Hsd17b4, Scarb2*), anti-angiogenesis (e.g. *Ctdsp1, Lyl1, Mapk14, Patz1, Ywhab*), NF-κB inhibitor (e.g. *Atp2a3, Bag2*), and pro-M2 (e.g. *Fads1, Klf2, Lpcat3, Pon2, Ywhab*) genes. Representative modulated genes are shown in Fig. 5C. A list of notable genes, along with annotations and statistical significances, is shown in Table S1. A complete list of unique 65-79*SE-modulated DEGs is shown in data file S2A.

**Fig. 5.**
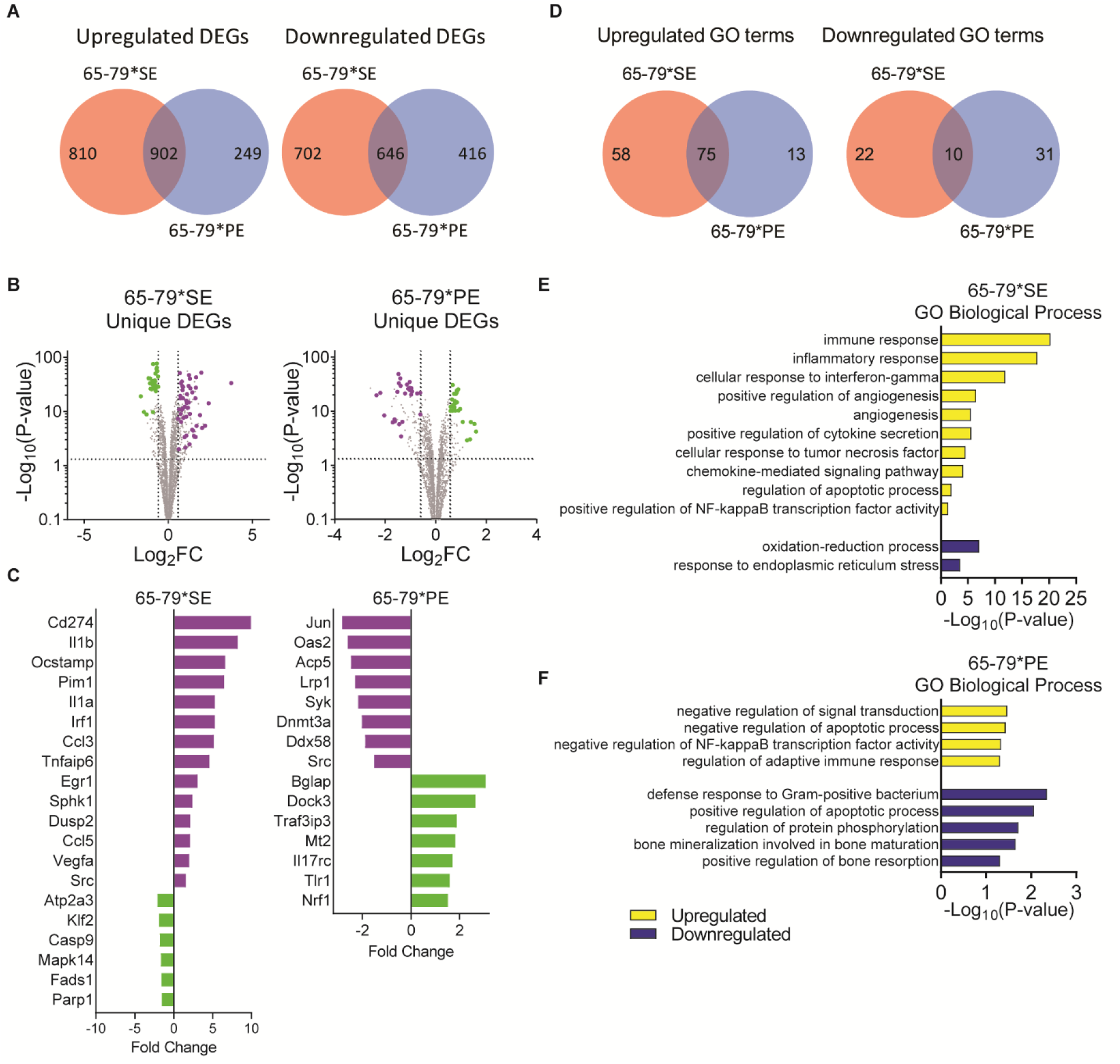
Transcription modulation by 65-79*SE and 65-79*PE under M1 polarizing conditions. Mouse RAW 264.7 macrophages were incubated for 3 days with IFNγ (5 ng/ml) in the presence or absence of 100 μg/ml 65-79*SE or 65-79*PE and RNA-seq analysis was performed on isolated RNA. Data are from 6 biological replicates in 2 independent experiments. (**A**) Venn diagrams showing upregulated and downregulated DEGs (P adjusted <0.05, fold change >1.5). (**B**) Volcano plots for unique DEGs for 65-79*SE or 65-79*PE. Purple dots denote pro-RA-associated genes; green dots denote RA-protective genes. (**C**) Selected notable genes unique for 65-79*SE or 65-79*PE with known RA-related functions. Purple bars denote pro-RA-associated genes; green bars denote RA-protective genes. (**D**) Venn diagrams of overlapping and unique GO terms derived from upregulated and downregulated DEGs in A. (**E, F**) Selected GO Biologic Processes for up- and downregulated DEGs by 65-79*SE (E) or 65-79*PE (F).

A diametrically opposite gene transcription pattern under M1-polarizing conditions was found with 65-79*PE (Fig. 5B and data file S2B). It showed uniquely upregulated anti-inflammatory (e.g. *Mt2, Il17rc, Havcr2, Ogg1, Nrf1, Stk10, Pdcd2*), anti-angiogenesis (e.g. *Flip1l, Fyn, Htatip2*), anti-bone resorption (e.g. *Def6, Gpr65, Bglap, Fbxl12*), anti-oxidative (e.g. *Mt2, Pycr1, Stc2, Dock3*) and pro-M2 or anti-M1 (e.g. *Tlr1, Nrf1, Themis2*) genes. Conversely, unique downregulated genes by 65-79*PE included many pro-osteoclastogenic (e.g. *Atp6v0d2, Dnmt3a, Syk, Ckb, Tspan5, Acp5, Src*), pro-angiogenesis (e.g. *St3gal1, Glul, Epn2, F7, Arhgap24*), pro-arthritogenic (e.g. *Syk*, *Jun*, *Tnfrsf9, F10, Arhgap24, Adamts7*), and NF-κB pathway-activating (e.g. *Sh3kbp1, Ddx58, F7)* genes. Representative genes are shown in Fig. 5C. A list of notable genes, along with annotations and statistical significances, is shown in Table S1. A complete list of unique 65-79*PE-modulated DEGs is shown in data file S2B.

Gene ontology (GO) analysis (Fig. 5D to F) revealed many GO processes that involve cytokine/chemokine signaling, innate immune response and positive regulation of NF-κB activity in macrophages stimulated with 65-79*SE under M1 polarizing conditions (Fig. 5E, data file S1C). By contrast, GO terms for genes upregulated by 65-79*PE in M1 polarizing conditions included processes involving inhibitory effects on signal transduction, protein phosphorylation, adaptive immune response and NF-κB (Fig. 5F, data file S1D). Conversely, GO terms for genes downregulated by 65-79*PE included bone mineralization and resorption, known as important effector mechanisms in RA pathogenesis. In summary, under M1 polarizing conditions, the PE-expressing TAHR activated an anti-inflammatory or anti-RA transcriptome, whereas the SE-expressing TAHR activated a pro-inflammatory, pro-RA transcriptome.

To determine whether TAHR polarizing effects could be found in human cells as well, we performed RNA-seq analysis in human THP-1 macrophages under M1 polarizing conditions, and found remarkable similarities between the two species (Fig. S4 and data file S3). For example, 100 (22%) of the 446 DEGs upregulated by 65-79*SE in human THP-1 cells were also upregulated by this TAHR in mouse RAW 264.7 macrophages (Fig. S4A, data file S3, A and B). Moreover, the top-ranked 60 DEGs that were found to be upregulated by 65-79*SE in both species were searched in PubMed, and 68% of them were found to be positively associated with RA disease risk or pathogenesis (Fig. S4B, Table S1 and data file S3B). GO analysis revealed many shared biologic processes (Fig. S4C, data file S3C). Moreover, KEGG pathways analysis (Fig. S4D, data file S3C) revealed a high level of correspondence between mouse and human macrophages in many relevant pathways, and identified the KEGG term Rheumatoid Arthritis as the top-ranked pathway in the THP-1 list.

A markedly different transcriptional landscape was observed under M2-polarizing conditions (Fig. 6 and data file S4). Notable amongst the unique upregulated genes by 65-79*PE in M2-polarizing conditions (Fig. 6C) were anti-angiogenic (*Glrx*, *Col7a1*, *Rras*), anti-inflammatory (*Anxa1, Rgs2, Tnfrsf17*), anti-bone remodeling (*Ctss*), NF-κB inhibitor (*Anxa1, Spn*), and activator of the Pi3K-Akt pathway (*Rras*) genes. Downregulated genes in these conditions included many pro-RA genes such as *Tnf*, *Fcrl1*, *Cxcl10*, *Il21r*, *Cxcl2*, *Ifi44l*, *Myo1d*, among others (Figs. 6A-6C, Table S1 and data file S4A). In spite of the anti-inflammatory effects by IL-4^42, 43^, 65-79*SE was able to upregulate various RA and pro-inflammatory genes (e.g. *Adamtsl5, Pyr1*, *Mmp9*, *Cd74*, *Cd84* and *Lat*) and downregulate several anti-RA, anti-inflammatory, or pro-M2 genes (e.g. *Rnase4, Col18a1Stk17b, Cd276, Cx3cr1*), although these 65-79*SE effects were weaker than its effect in M1-polarizing conditions (Fig. 6A-6C, Table S1 and data file S4B). GO analysis (Figs. 6D-6F) showed that pathways related to immunity and inflammation were upregulated by 65-79*SE and downregulated by 65-79*PE (Fig. 6E and F, data files S4C and D). Thus, under M2 polarizing conditions 65-79*PE and 65-79*SE had reciprocal effects; the former enhanced anti-inflammatory effects, whereas the latter was capable of moderately enhancing an inflammatory transcription profile.

**Fig. 6.**
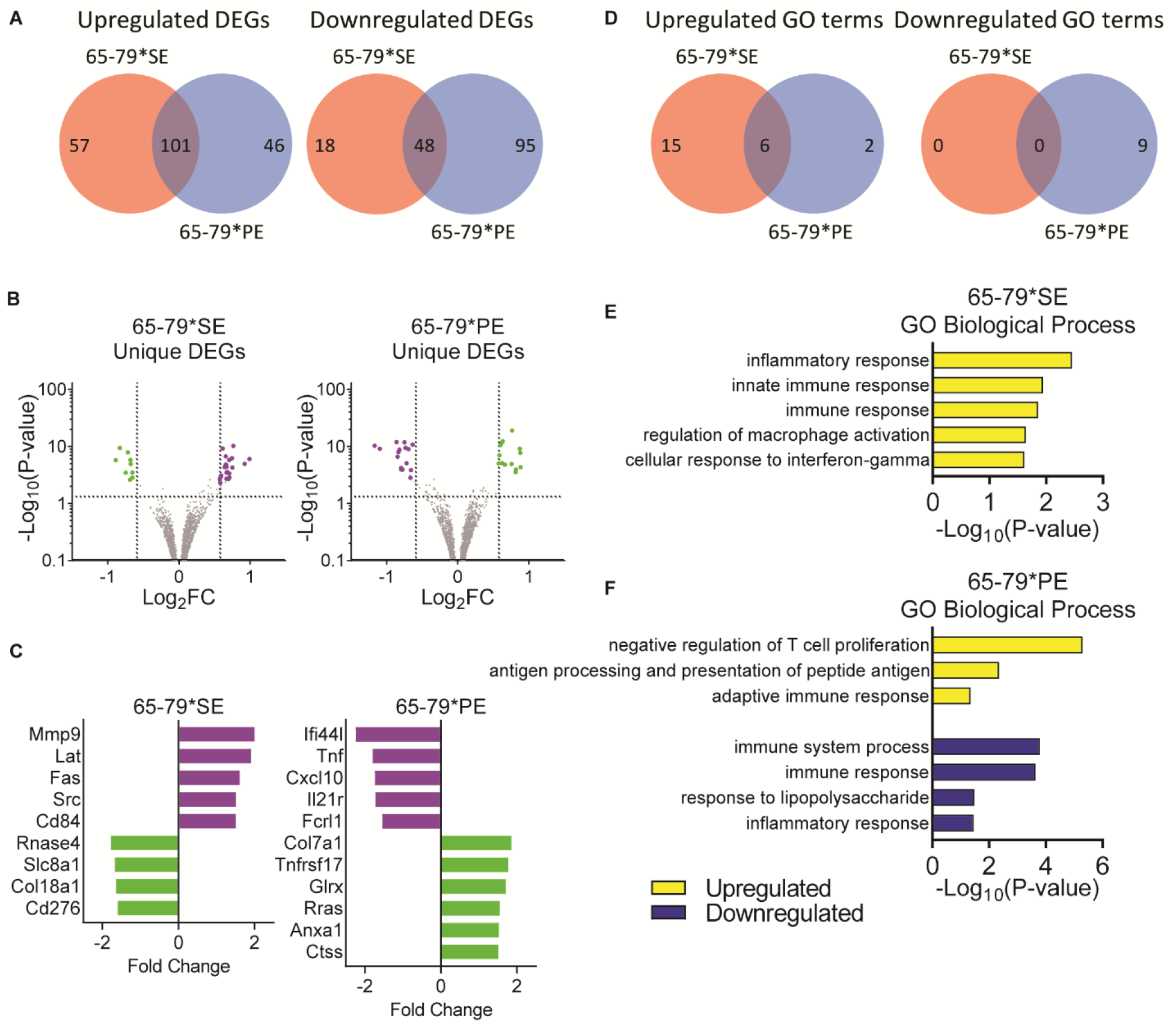
Transcription modulation by 65-79*SE and 65-79*PE under M2 polarizing conditions. Mouse RAW 264.7 macrophages were incubated for 3 days with Il-4 (5 ng/ml) in the presence or absence of 65-79*SE or 65-79*PE and RNA-seq analysis was performed as in Fig. 5. Data are from 6 biological replicates in 2 independent experiments. (**A**) Venn diagrams showing upregulated and downregulated DEGs (P adjusted <0.05, fold change >1.5). (**B**) Volcano plots for unique DEGs for 65-79*SE or 65-79*PE. Purple dots denote pro-RA-associated genes; green dots denote RA-protective genes. (**C**) Selected notable genes unique for 65-79*SE or 65-79*PE with known RA-related functions. Purple bars denote pro-RA-associated genes; green bars denote RA-protective genes. (**D**) Venn diagrams of overlapping and unique GO terms derived from upregulated and downregulated DEGs in A. **(E**, **F**) Selected GO Biologic Processes for up- and downregulated DEGs by 65-79*SE (E) or 65-79*PE (F).

### Upstream Regulators

To assess the underlying mechanisms driving the gene expression patterns that are induced by 65-79*SE and 65-79*PE, we performed upstream regulator analyses. In M1-polarizing conditions (Fig. 7A-C, Table S2 and data file S5A), 65-79*SE was predicted to stimulate activation of pro-inflammatory, pro-RA upstream regulators, such as Stat3, Rel, Rela, Jun, Ctnnb1 and Hif1a, among others, and to inhibit anti-arthritis or anti-inflammatory regulators (e.g. Klf2, Xbp1, Foxp3) (Fig. 7A). In these conditions, 65-79*PE was predicted to activate several anti-inflammatory or NF-κB-inhibiting regulators, such as Nfkbiz, Mta1, Tardbp, Xbp1, and inhibit several key transcription factors known to associate with osteoclastogenesis (Mitf, E2f1), angiogenesis (Srebf1, E2f1, Sox11) and inflammation (Srebf1, Keap1), among others (Fig. 7B, Table S2 and data file S5B).

**Fig. 7.**
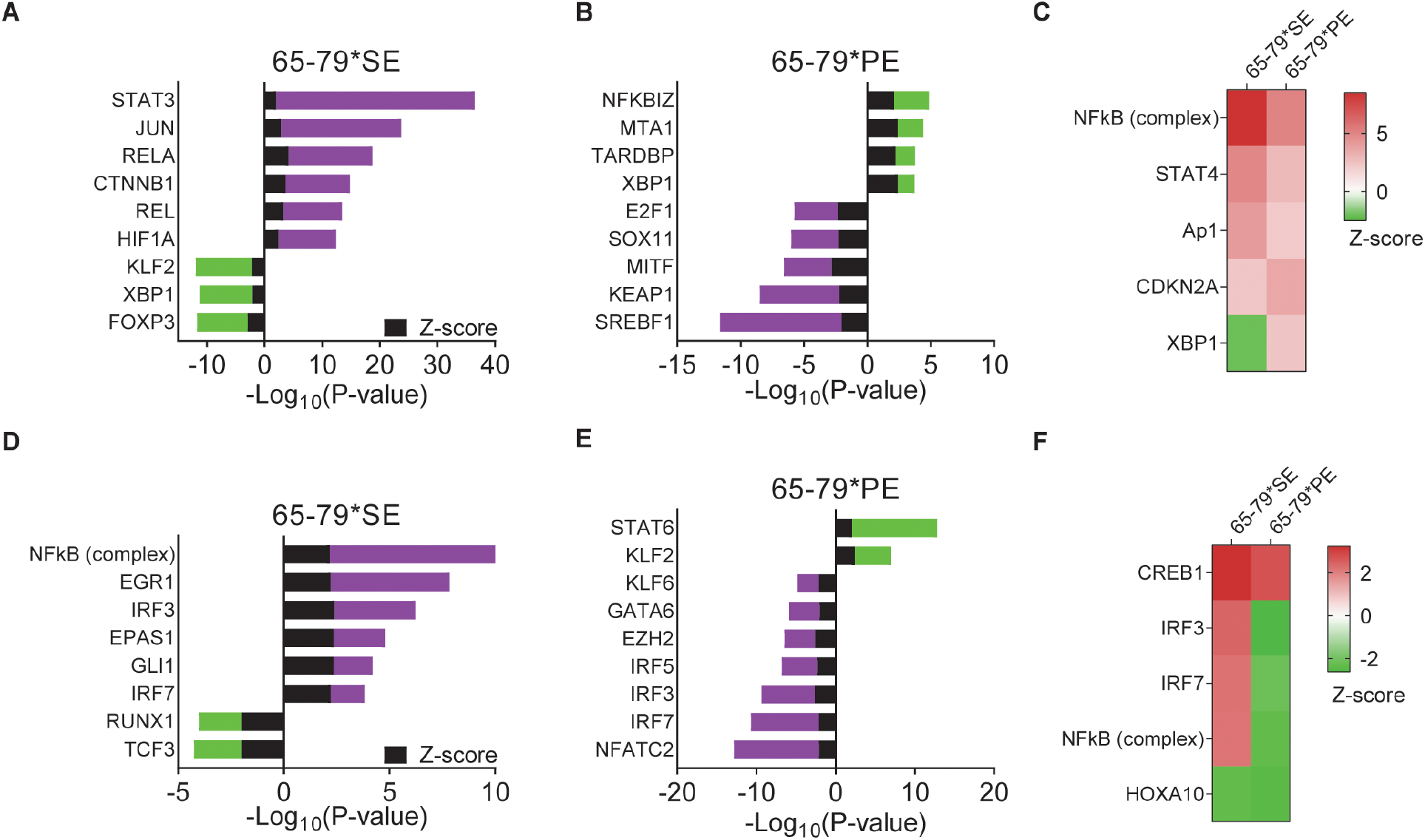
Differential activation of upstream regulators by 65-79*SE and 65-79*PE under M1 or M2 polarization conditions. (**A**, **B**) Predicted activated and inhibited upstream regulators by 65-79*SE (A), 65-79*PE (B) in M1-polarizing conditions. (**C**) Heatmap of notable upstream regulators for 65-79*SE and 65-79*PE in M1 polarizing conditions. (**D**, **E**) Predicted activated and inhibited upstream regulators by 65-79*SE (C), or 65-79*PE (D) in M2-polarizing conditions. In (A, B, D, E); purple bars denote pro-inflammatory or pro-RA regulators; green bars indicate anti-inflammatory or anti-RA regulators. (**F**) Heatmap of notable upstream regulators for 65-79*SE and 65-79*PE in M2 polarizing conditions. In (C) and (F); red denotes predicted activation; green denote predicted inhibition.

Under M2 polarizing conditions (Fig. 7D-F, Table S2 and data file S5, C and D), 65-79*SE was predicted to exert an activation effect on several pro-inflammatory or pro-angiogenic upstream regulators, including Egr1, Irf3, Irf7, Epas1, Gli1 and NF-κB, and an inhibitory effect on anti-RA, anti-osteoclastogenic or anti-angiogenic upstream regulators, such as Tcf3 and Runx1 (Fig. 7D). Conversely, 65-79*PE was predicted to activate key M2-inducing and anti-arthritis transcription factors Stat6 and Klf2, and inhibit pro-arthritis (Irf5, Irf7), pro-osteoclastogenic (Nfatc2, Ezh2), pro-angiogenic (Nfatc2, Ezh2, Gata6, Klf6, Irf3), pro-inflammatory (Irf5, Klf6), and pro-M1 (Klf6, Irf3, Irf7) upstream regulators (Fig. 7E). Intriguingly, several upstream regulators were predicted to be modulated in diametrically opposite directions by 65-79*SE versus 65-79*PE. For example, in M1-polarizing conditions (Fig. 7C), Xbp1, an anti-inflammatory and NF-κB-inhibiting upstream regulator, was predicted to be inhibited by 65-79*SE, yet activated by 65-79*PE. In M2-polarizing conditions (Fig. 7F), a pro-M1 and pro-angiogenic upstream regulator Irf3, and the pro-M1 pro-arthritis upstream regulator Irf7, and NF-κB were all predicted to be activated by 65-79*SE, and reciprocally inhibited by 65-79*PE. Importantly, consistent with the above findings, and the known role of NF-κB in the pathogenesis of autoimmune diseases, upstream regulators analysis confirmed a pivotal role for NF-κB complex pathway, with opposite outcomes in the presence of 65-79*SE versus 65-79*PE (Fig. 8).

**Fig. 8.**
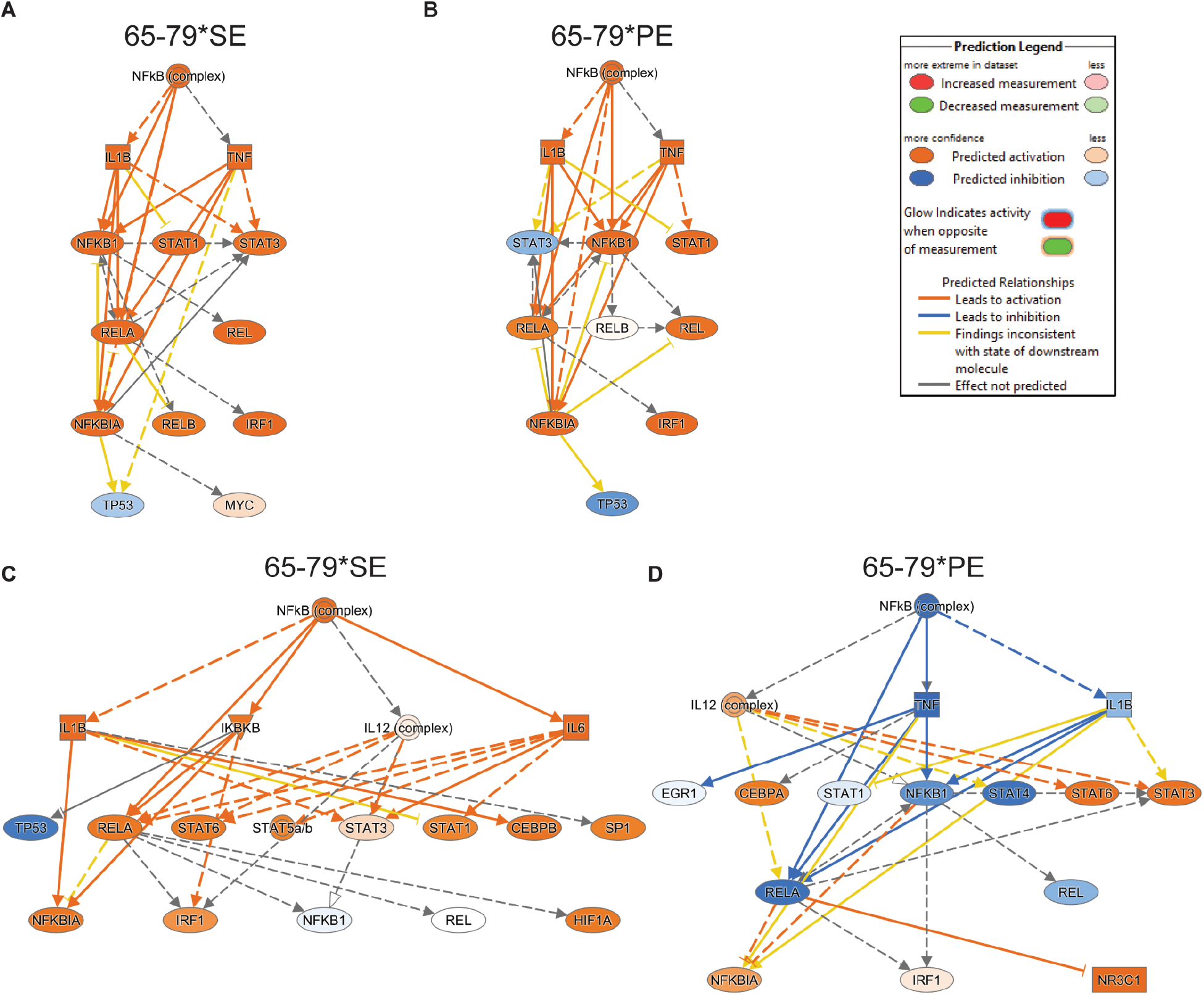
Differential NF-κB pathway activation by 65-79*SE and 65-79*PE under M1 polarizing conditions (A, B) or under M2 polarization conditions (C, D)

## Discussion

Decades after MHC-restricted AP^2, 3^ and HLA-disease association (reviewed in^1^) were independently discovered, it remains unclear whether the two processes are mechanistically-related. Evidence suggesting an antigen-specific immune response exists in some HLA-associated diseases, but in the majority of conditions, a candidate target antigen has not been identified. The findings of this study implicate an allele-specific AP-independent mechanism.

Given on the known pivotal role that macrophages play in regulating pro- and anti-inflammatory events, here we focused on these cells. Our findings revealed differential activation of macrophage polarization pathways by *HLA-DRB1* allele-specific gene products. Using primary macrophages derived from two transgenic mouse lines that express distinct human HLA-DRβ molecules, which differ by only 3 amino acid residues in the TAHR, we observed diametrically opposite polarization patterns. Primary macrophages derived from SE Tg (transgenic mice expressing the SE motif 70-QKRAA-74 in the TAHR of the DRβ chain) showed strong predilection to pro-inflammatory M1 macrophage differentiation, while cells derived from PE Tg (expressing a 70-DERAA-74 sequence), displayed preferential differentiation of M2-type macrophages.

The two alleles differentially regulated signaling events. Under M1-polarizing culture conditions, primary macrophages from PE Tg were found to display increased Akt activation and attenuated M1 polarization. In macrophages from SE Tg, on the other hand, SHIP1-mediated Akt inhibition was implicated in enhanced M1 polarization. Under M2-polarizing conditions PE Tg macrophages, but not SE Tg macrophages, selectively enhanced M2 gene expression via the Akt - and possibly Stat6 - pathways. Thus, autoimmune disease susceptibility or -protective *HLA-DRB1* alleles were found to reciprocally regulate signaling pathways, which determined the efficiency of pro-inflammatory versus anti-inflammatory macrophage differentiation outcomes.

*In vitro-*differentiated BMDM cultures are devoid of lymphocytes. It is therefore unlikely that the differential macrophage polarization observed here involved AP. To ascertain that AP is indeed not involved, and to determine whether the effect could be mapped to the TAHR, we used AP-incompetent 15-mer peptides that correspond to residues 65-79 in the DRβ chain to explore their transcriptional activation in mouse RAW 264.7 and human THP-1 macrophages. The findings revealed that: A. Synthetic peptides corresponding to TAHRs that differ by only 3 amino acid residues activated allele-specific signature transcriptomes; B. The effect was AP-independent, since it was activated by short, AP-incompetent synthetic peptides; C. RNA-seq parallels between mouse and human macrophages strengthen the significance of the findings.

Under M1-polarizing conditions, 65-79*SE upregulated the expression levels of many genes that are known to code for pro-inflammatory or known RA disease markers, as well as genes coding for confirmed - or proposed - therapeutic targets, and pathogenic mechanisms, such as osteoclastogenesis, NF-κB activation, angiogenesis, or M1 polarization. Conversely, the SE-expressing TAHR downregulated anti-inflammatory, anti-angiogenesis, inhibitors of NF-κB, and pro-M2 genes. 65-79*PE, on the other hand, upregulated many anti-inflammatory, pro-M2 genes, as well as RA-protective, anti-oxidant, genes. Under M2 polarizing conditions, 65-79*PE downregulated genes and GO processes that are associated with inflammation and RA pathogenesis, while 65-79*SE, notwithstanding the anti-inflammatory tissue culture milieu, managed to upregulate some genes and GO processes known to mediate pro-inflammatory and macrophage activation events.

Importantly, upstream regulator analysis based on gene expression patterns predicted the NF-κB-mediated pathway to be activated by the SE and inhibited by the PE (Fig. 8). This finding is congruent with the key role that NF-κB plays in autoimmune diseases^44, 45, 46, 47, 48, 49^. Thus, consistent with their reciprocal impacts on autoimmune disease protection versus susceptibility, the two epitopes dampen (PE) or enhance (SE) pro-inflammatory events.

An RA-protective effect of 70-DERAA-74-coding *HLA-DRB1* alleles has been known for some time^21^, but the mechanism underlying this effect has been unclear. A recent study proposed that antigenic mimicry between vinculin and bacterial proteins presented by HLA-DQ molecules, which are commonly associated with SE-coding *HLA-DRB1* alleles through linkage disequilibrium may be involved in RA^50^. However, that hypothesis is inconsistent with the fact that PE-coding *HLA-DRB1* alleles have a protective effect in SE-negative individuals as well^51^. Moreover, as discussed above, in addition to their protective effect in RA, PE-coding *HLA-DRB1* alleles have been found to decrease disease risk in many other autoimmune conditions that do not share putative antigens with RA. We propose that the AP-independent mechanism identified here is more plausible.

Another group reported recently that in contrast to their protective effect when carried through Mendelian inheritance, DERAA-coding *HLA-DRB1* alleles increase RA disease susceptibility when acquired through pregnancy-associated microchimerism^52^. That study reported that the microchimerism was prevalence mostly in early RA, with diminished prevalence in patients with chronic stages of the disease^52^. The mechanisms underlying the dichotomous effect of PE-coding *HLA-DRB1* alleles, and whether AP is involved remain unknown. Given the diminishing prevalence over time, it is tempting to speculate that the phenomenon may be a consequence of initial gestational tolerance to the allogeneic PE, followed by post-delivery “rebound” of an immune response that leads to clearance of the alloantigen, but due to immune cross reactivity against a putative non-*HLA-DRB1*-coded disease-protective factor, a paradoxical increased risk of RA is found in these patients.

It is also worth mentioning that based on imputation of genomics data it has been suggested by others that in addition to the 5 residues 70-74 in the TAHR that determine the SE-associated RA disease risk, peptide-binding groove residues 11 and 13 associate significantly with RA risk as well^53^, indirectly suggesting that AP may be involved. However, this imputation-based theory has not yet been experimentally validated, and the relevance of the observation to RA pathogenesis has been questioned^54^. Be that as it may, the mechanism proposed here does not exclude possible involvement of AP in HLA-associated diseases; it offers explanation to aspects of HLA-disease associations that are inconsistent with AP alone. It is not inconceivable that while presentation of specific antigen(s) may determine the anatomic site(s) involved, the SE and PE polarize the immune response to pro-versus anti-inflammatory modes, thereby shaping the pathogenic outcomes.

In summary, our findings lend support to the MHC Cusp theory^8, 9^, which posits that independent of their known role in AP, MHC molecules express allele-specific epitopes which can activate cell signaling events that play a role in HLA-disease associations. Extending this line of pursuit to other alleles and their associated conditions could determine the broader relevance of these findings.

## Materials and Methods

### Mice

Transgenic mice, expressing the human *HLA-DR4*04:01* or *HLA-DR4*04:02* alleles^55, 56^ were kindly provided by Dr. Chela David, at the Mayo Clinic, and are referred to as SE Tg and PE Tg, respectively. The two mouse strains have a mixed (predominantly B6) genetic background and are approximately 99% identical. 10-12 week old male mice were housed under specific pathogen-free and temperature-controlled (25°C) conditions in a 12-hr dark/light cycle. All experimental mouse protocols were approved by the University of Michigan Unit for Laboratory Animal Medicine and by the University of Michigan Committee on Use and Care of Animals. All applicable federal, state, local, and institutional laws, regulations, policies, and standards governing animal research were followed. In some experiments, LPS (500 μg/kg) was administered to 10-12 week old male mice using a single i.p. injection. Serum or peritoneal exudate cells (PECs) were collected 4 hrs. after i.p. injection.

### Reagents

All reagents used, along with vendor names and catalog numbers are listed in supplementary Table S3.

### Primary Macrophage Culture Conditions

Primary mouse BMDMs were isolated and cultured as previously described^57^. Peritoneal macrophages were isolated by injecting ice-cold PBS into the peritoneal cavity. PECs were subsequently collected, centrifuged and washed once with PBS. For qRT-PCR experiments, macrophages were cultured for 3 days in 6-well plates (2 × 10^6^ cells per well) in α-MEM with 10% (v/v) FBS, 100 U/ml penicillin and 100 μg/ml streptomycin, along with 10 ng/ml recombinant macrophage colony stimulating factor (M-CSF). Culture media were refreshed daily. In cell function experiments, instead of M-CSF, macrophages were cultured for 3 days in 20% (v/v) L929 cell conditioned media, in addition to 0.5% (v/v) pyruvate, 10% (v/v) FBS, 100 μg/ml streptomycin and 100 U/ml penicillin. To induce M1 polarization, cells were treated with 1 ng/ml LPS and 20 ng/ml IFNγ for 24 hrs. To induce M2 polarization, Il-4 and Il-13 (10 ng/ml each) were added for 24 hrs. For experiments with inhibitors, cells were treated with Ly294004 (5 μM), PF-4708671 (10 μM), or wedelolactone (10 μM) for 1hr prior to polarization.

### Macrophage Cell Lines Culture Conditions

Mouse RAW 264.7 macrophages were maintained in DMEM containing 10% (v/v) FBS, 100 μg/ml streptomycin and 100 U/ml penicillin. Cells were cultured in T75 flasks to confluence and were split every 3 days. THP-1 cells were cultured in a 10% (v/v) FBS-RPMI 1640 medium containing 100 U/ml penicillin and 100 μg/ml streptomycin. To differentiate THP-1 cells into macrophages they were cultured for 3 days with phorbol 12-myristate 13-acetate (PMA, 85 nM). Differentiated cells were then cultured without PMA for an additional 5 days prior to the experiments.

### RNA isolation and qRT-PCR

Mouse cells were lysed with Trizol for RNA isolation. Direct-Zol™ RNA miniprep (Zymo research) was used to isolate total RNA. Isolation of RNA from THP-1 cells was carried out with a RNeasy Plus Mini kit (Qiagen). The High-Capacity cDNA Reverse Transcription Kit (Applied Biosystems) was used to synthesize cDNA. qRT-PCR was performed by Fast SYBRTM Green Master Mix (Applied Biosystems), with sets of primers as listed in supplementary Table S4, using a StepOnePlus Real-Time PCR system (Applied Biosystems). A StepOne Software was used to analyze the data with the ΔΔCT method.

### Immunoblots

After being washed with ice-cold PBS, cells were lysed in RIPA buffer (Sigma) with EDTA-free protease phosphatase inhibitors (Roche Diagnostics). Protein concentration in lysates was determined using the DC protein assay (BioRad). Proteins were loaded onto 4-20% SDS–PAGE gels (Invitrogen), and after electrophoretic separation, transferred to nitrocellulose membranes (BioRad), followed by incubation with appropriate primary and secondary antibodies. All primary antibodies were diluted 1:1000 in 5% (v/v) BSA (Sigma-Aldrich), except β-Actin (1:4,000). The primary antibodies used were anti: Akt (#9272), pAkt (S473, #9271), STAT6 (#9362), pSTAT6 (Y641, #56554), SHIP1 (#2728), pSHIP1 (Y1020, #3941), PTEN (#9559), pPTEN (S380/T382/383, #9549) (all from Cell Signaling Technology), or anti-β-Actin (Invitrogen, BA3R). Second-stage antibodies included Anti-Rabbit IgG HRP-conjugated (Cell Signaling Technology, 1:1000) or Anti-mouse IgG HRP-conjugated (GE Healthcare, 1:8000). Proteins visualization was performed using a SuperSignal West Pico Plus ECL substrate (Thermo Scientific) and an Omega Lum C imaging system (Gel Company). Band quantification was performed in duplicates with the ImageJ software.

### Cytokine measurements

Tnfα, Il-12p70, Il-10 and Il-6 were quantified in culture supernatants by ELISA kits (R&D). Serum cytokine levels were measured using Quantibody Mouse Cytokine array 1 (RayBiotech). Slides were scanned by RayBiotech Service Department and data were analyzed using a RayBiotech Mouse Cytokine Array 1 software (QAM-CYT-1-SW). Sample concentrations were derived from mean fluorescence intensities, relative to standard curves, generated using the manufacturer’s standards.

### NO and Arginase assays

NO production was quantified using the fluorescent NO dye 4,5-diaminofluorescein diacetate (DAF-2DA) as previously described^58^. Cells were first plated overnight in flat-bottom, 96-well plates, then washed with DMEM/phenol red–free medium (Sigma). Cultures were then loaded with 20 μM DAF-2DA at 37°C for 1 hr. in the dark. Fluorescence levels were recorded every 5 minutes over a period of 500 minutes using a Synergy H1 hybrid reader system (Biotek) at excitation/emission wavelengths of 488/515 nm. NO production rates are expressed as fluorescence units (FU) per minute. Measurement of arginase activity was carried out using the Arginase Activity Colorimetric Assay Kit (Biovision).

### RNA-Seq

RAW 264.7 cells were incubated with or without IFNγ (5 ng/ml) or Il-4 (5 ng/ml) in the presence or absence of 15-mer peptides 65-79*SE or 65-79*PE (100 μg/ml) and medium was refreshed at 48 hrs. At 72 hrs., total RNA was isolated using the Direct-Zol™ RNA miniprep (Zymo research). Genomic DNA was removed using the Turbo DNA-Free kit (Ambion). Total RNA concentration and integrity were determined with an Agilent Bioanalyzer (Agilent). All RNA samples had an integrity number of 7.5 or higher. Two hundred ng of total RNA were used to generate libraries using the TruSeq Stranded mRNA Sample Preparation Kit (Illumina). Single-end reads of 100bp for each sample were produced with Illumina’s HiSeq2500v4 instrument. Raw counts were attained using featureCounts from the Rsubread1.5.0p3 package and Gencode-M12 gene annotations using only uniquely aligned reads. For THP-1, Raw counts were attained using featureCounts from the Rsubread-1.6.1 package and - Gencode28-hg38 gene annotations using only uniquely aligned reads. DESeq2-1.16.1 within R-3.4.1 was used to perform data normalization and differential expression analysis with an adjusted p-value threshold of 0.05. Mouse annotations were verified using the MGI database (http://www.informatics.jax.org/index.shtml)^59^ and only protein encoding genes were considered. Genes with a fold-change of >1.5 and p-values of 0.05 were used for GO term analysis. RA relevant genes were identified based on published literature. The DAVID bioinformatics database (version 6.8)^60^ was used for GO analysis, using an Expression Analysis Systematic Explorer (EASE) score threshold of 0.1 for detection of gene enrichment. GO terms with an FDR of 5% were considered significant. To identify unique GO terms, significant GO terms obtained in 65-79*SE-stimulated cells in either M1- or M2-polarizing culture conditions were compared to those obtained in 65-79*PE-stimulated cells under the same polarizing culture conditions.

### Upstream Regulator Analysis

Ingenuity Pathway Analysis (Qiagen) was used to perform upstream regulator analysis. Genes differentially expressed in the presence of 65-79*SE or 65-79*PE were analyzed to identify transcriptional regulators that may be responsible for the gene expression changes observed. Only transcriptional regulator relationships observed using experimental data were used in this analysis. A z-score was used to examine the robustness of predicted activation or inhibition of specific transcription regulators based on enrichment and differential expression (upregulation or downregulation) in the presence of 65-79*SE or 65-79*PE. P values <0.05 by Fisher’s Exact Test after correction for multiple testing were considered significant.

### Statistics

Data are shown as the mean and SEM calculated using a GraphPad Prism Software (version 7). Unless otherwise specified, a 2-way ANOVA was used to determine significance. P-values < 0.05 were considered significant.

## Supplementary Materials

Fig. S1. Role of Pi3K regulating phosphatases.

Fig. S2. Differential Stat6 phosphorylation in SE Tg and PE Tg BMDMs under M2 polarizing conditions.

Fig. S3. Differential transcriptional response to 65-79*SE or 65-79*PE under M1 and M2 polarizing conditions.

Fig. S4. Corresponding upregulated DEGs between THP-1 and RAW 264.7 cells.

Table S1. Notable DEGs with RA relevance.

Table S2. Notable upstream regulators with RA relevance.

Table S3. List of reagents used in this study.

Table S4. List of primer sequences used for qPCR analysis.

Data file S1 (Microsoft Excel format). RNA-seq data (RAW 264.7 cells): All DEGs and GO analysis under M1 polarization conditions.

Data file S2 (Microsoft Excel format). RNA-seq data (RAW 264.7 cells): Unique DEGs under M1 or M2 polarization conditions.

Data file S3 (Microsoft Excel format). RNA-seq data (THP-1 cells): DEGs and GO terms under M1 polarization conditions.

Data file S4 (Microsoft Excel format). RNA-seq data (RAW 264.7 cells): All DEGs and GO analysis under M2 polarization conditions

Data file S5 (Microsoft Excel format). Upstream regulators under M1 or M2 polarization conditions

## Acknowledgments

We thank E. Henson, T. Tubo, Y. Liu and Dr. W. H. Ali-Hanel at the University of Michigan for technical assistance, and Dr. P. Gourh and Dr. P. Grayson at the National Institute of Arthritis and Musculoskeletal and Skin Diseases for useful comments on the manuscript.

## Funding

This work was supported by the Extramural Program of the National Institute of Arthritis and Musculoskeletal and Skin Diseases (Grants R01AR059085, R61AR073014, R33AR073014, R01ARR074930) to J.H., the National Institute of Environmental Health Sciences Extramural Program (Contract HHSN273201600123P) to J.H., and the Intramural Research Program at the National Institute of Environmental Health Sciences (Grant ES101074) to F.W.M. The content is solely the responsibility of the authors and does not necessarily represent the official views of the National Institutes of Health.

## Author contributions

J.H. conceived the study. V.vD., B.M.S., A.H.S. and J.H. designed the experiments. V.vD., S.V.N., and B.M.S. performed the experiments. V.vD., B.M.S., A.H.S. and J.H. analyzed data. F.W.M. and J.H. provided funding. V.vD. and J.H. wrote the paper. All authors interpreted the data, reviewed the paper, and approved the final version of the manuscript.

## Competing interests

J.H. is an Inventor of Regents of the University of Michigan-owned technologies that are licensed to Zydus-Caldia, to whom he is an unpaid consultant, or Alibion AG, to whom he is a paid consultant. All other authors have no competing interests to declare.

## Data and materials availability

The RNA-seq data are deposited on GEO database (GEO number: GSEXXX).

