## Supplemental Materials for "Antigen Presentation-Independent Reciprocal Immune Modulation by *HLA-DRB1* Allelic Epitopes that Associate with Autoimmune Disease Risk or Protection"

**The PDF file includes:**

Fig. S1. Role of Pi3K regulating phosphatases.

Fig. S2. Differential Stat6 phosphorylation in SE Tg and PE Tg BMDMs under M2 polarizing conditions.

Fig. S3. Differential transcriptional response to 65-79*SE or 65-79*PE under M1 and M2 polarizing conditions.

Fig. S4. Corresponding upregulated DEGs between THP-1 and RAW264.7 cells.

Table S1. Notable DEGs with autoimmune disease relevance.

Table S2. Notable upstream regulators with autoimmune disease relevance.

Table S3. List of reagents used in this study.

Table S4. List of primer sequences used for qPCR analysis.

**Other Supplemental Materials include:**

Data file S1 (Microsoft Excel format). RNA-seq data (RAW 264.7 cells): All DEGs and GO analysis under M1 polarization conditions

Data file S2 (Microsoft Excel format). RNA-seq data (RAW 264.7 cells): Unique DEGs under M1 or M2 polarization conditions

Data file S3 (Microsoft Excel format). RNA-seq data (THP-1 cells): DEGs and GO terms under M1 polarization conditions

Data file S4 (Microsoft Excel format). RNA-seq data (RAW 264.7 cells): All DEGs and GO analysis under M2 polarization conditions

Data file S5 (Microsoft Excel format). Upstream regulators under M1 or M2 polarization conditions

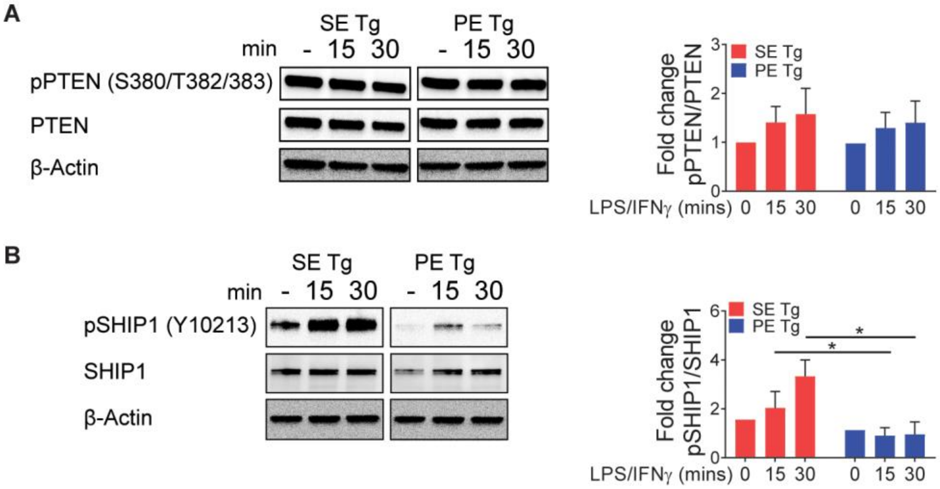

**Fig. S1. Role of Pi3K regulating phosphatases**

(**A**) Immunoblot for pPTEN (Ser380/Thr382/383) and PTEN under M1 polarizing conditions [LPS (1 ng/ml) + IFNγ (20 ng/ml)] for 15-30 min.

(**B**) Immunoblot for pSHIP1 (Tyr1021) and SHIP1 under M1 polarizing conditions as in (A).

Quantification data represent mean + SEM of 3 independent experiments. 2-way ANOVA, *$p<0.05$

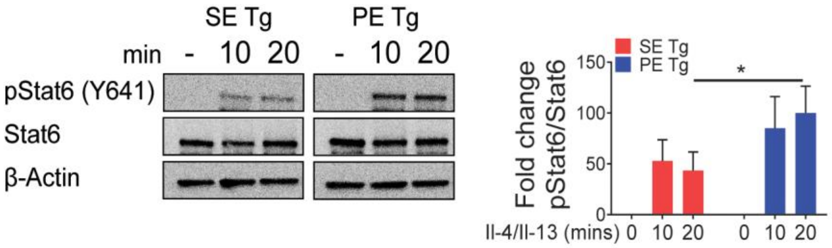

**Fig. S2. Differential Stat6 phosphorylation in SE Tg and PE Tg BMDMs under M2 polarizing conditions**

Western blot of pStat6 (Tyr641) and Stat6 in BMDMs derived from SE Tg or PE Tg mice following treatment with IL-4 (10 ng/ml) and IL-13 (10 ng/ml) for 10 or 20 minutes. Data on the right represent mean and SEM of 3 independent experiments. 2-way ANOVA, *P<0.05.

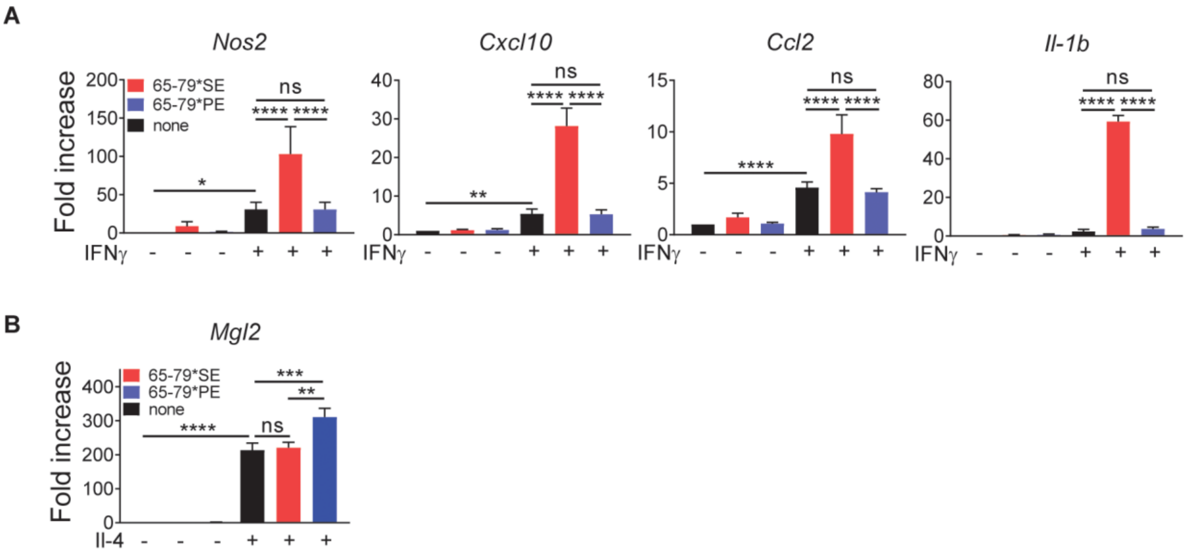

**Fig. S3**. **Differential transcriptional response to 65-79*SE or 65-79*PE under M1 and M2 polarizing conditions**

Mouse RAW 264.7 macrophages were incubated for 3 days with IFNγ (5 ng/ml) or IL-4 (5 ng/ml) in the presence or absence of 100 µg/ml 65-79*SE or 65-79*PE.

(**A**) qRT-PCR for the M1 genes *Nos2*, *Cxcl10*, *Ccl2* and *Il-1b*. Data represent mean + SEM of 3 biological replicates.

(**B**) qPCR for M2 gene *Mgl2*. Data represents mean + SEM of 5 biological replicates, 1-way ANOVA, *P<0.05, **P<0.01, ****P<0.0001.

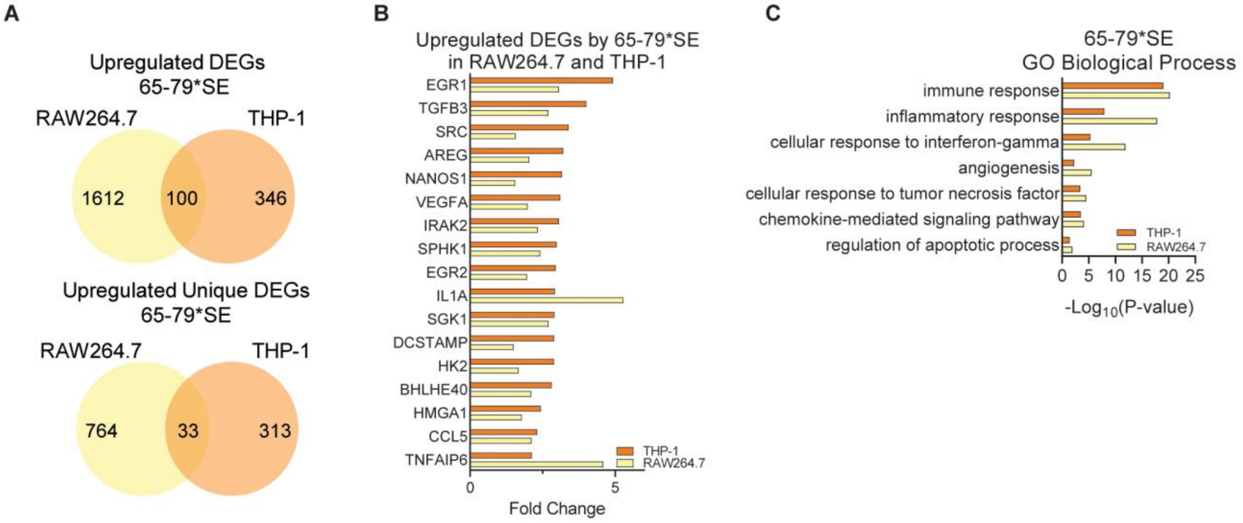

D

| | THP-1 | | RAW264.7 | |
|---|---|---|---|---|
| Term | Benjamini | Padj | Benjamini | Padj |
| hsa05323:Rheumatoid arthritis | 5.06E-17 | 5.06E-19 | 2.94E-07 | 2.94E-09 |
| hsa05332:Graft-versus-host disease | 2.51E-14 | 2.51E-16 | 2.08E-13 | 2.08E-15 |
| hsa04940:Type I diabetes mellitus | 2.01E-12 | 2.01E-14 | 2.41E-12 | 2.41E-14 |
| hsa04612:Antigen processing and presentation | 2.74E-12 | 2.74E-14 | 1.68E-13 | 1.68E-15 |
| hsa05330:Allograft rejection | 5.89E-11 | 5.89E-13 | 6.24E-12 | 6.24E-14 |
| hsa05416:Viral myocarditis | 2.32E-10 | 2.32E-12 | 7.81E-09 | 7.81E-11 |
| hsa05320:Autoimmune thyroid disease | 8.13E-09 | 8.13E-11 | 1.47E-08 | 1.47E-10 |
| hsa05140:Leishmaniasis | 6.46E-08 | 6.46E-10 | 3.42E-08 | 3.42E-10 |
| hsa05152:Tuberculosis | 7.57E-08 | 7.57E-10 | 3.08E-08 | 3.08E-10 |
| hsa05321:Inflammatory bowel disease (IBD) | 1.11E-07 | 1.11E-09 | 3.63E-08 | 3.63E-10 |
| hsa04514:Cell adhesion molecules (CAMs) | 9.81E-07 | 9.81E-09 | 7.41E-10 | 7.41E-12 |
| hsa05150:Staphylococcus aureus infection | 1.00E-06 | 1.00E-08 | 8.89E-14 | 8.89E-16 |
| hsa04145:Phagosome | 2.03E-06 | 2.03E-08 | 6.40E-12 | 6.40E-14 |
| hsa05168:Herpes simplex infection | 2.22E-06 | 2.22E-08 | 2.21E-12 | 2.21E-14 |
| hsa05310:Asthma | 2.40E-06 | 2.40E-08 | 1.82E-04 | 1.82E-06 |
| hsa05166:HTLV-I infection | 9.53E-06 | 9.53E-08 | 1.77E-12 | 1.77E-14 |
| hsa04060:Cytokine-cytokine receptor interaction | 1.35E-05 | 1.35E-07 | 8.24E-12 | 8.24E-14 |
| hsa05145:Toxoplasmosis | 1.36E-05 | 1.36E-07 | 1.93E-08 | 1.93E-10 |
| hsa04672:Intestinal immune network for IgA production | 1.37E-05 | 1.37E-07 | 6.94E-08 | 6.94E-10 |
| hsa05164:Influenza A | 8.04E-04 | 8.04E-06 | 1.49E-08 | 1.49E-10 |
| hsa05169:Epstein-Barr virus infection | 8.26E-04 | 8.26E-06 | 2.61E-05 | 2.61E-07 |
| hsa04668:TNF signaling pathway | 9.07E-04 | 9.07E-06 | 2.55E-05 | 2.55E-07 |

**Fig. S4. Corresponding upregulated DEGs between THP-1 and RAW 264.7 cells under M1 polarizing conditions**

THP-1 (human) or RAW 264.7 (mouse) macrophages were incubated for 3 days with IFNγ (5ng/ml) in the presence or absence of 100 μg/ml 65-79*SE and RNA-seq analysis was performed on isolated RNA. THP-1 data are from 4 independent experiments. RAW 264.7 data are from 6 biological replicates in 2 independent experiments.

(**A**) Venn diagrams showing the overlap between THP-1 and RAW 264.7 cells for all and unique upregulated DEGs (P adjusted <0.05, fold change >1.5).

(**B**) Notable unique genes with autoimmune disease relevance upregulated by 65-79*SE in both THP-1 and RAW 264.7 cells.

(**C**) Notable GO Biologic Processes unique for 65-79*SE for upregulated DEGs in both THP-1 and RAW 264.7 cells.

(**D**) KEGG pathways unique for 65-79*SE for upregulated DEGs in both THP-1 and RAW 264.7 cells.

### Table S1: DEGs with autoimmune disease relevance

#### S1A. Notable unique DEGs modulated by 65-79*SE under M1 polarizing conditions

| Upregulated | | | |
|---|---|---|---|
| | Fold Change | Padj | Relevant roles/functions |
| Dusp4 | 3.9137 | 1.61E-52 | Pro-angiogenic;<br>Pro-Th17 polarization |
| Vav1 | 1.6236 | 1.44E-51 | RA-associated;<br>Neutrophil activation;<br>Involved in CD28-mediated T cell activation through NF-kB pathway |
| Cd44 | 2.3622 | 1.95E-47 | MIF co-receptor; plays a role in RA pathogenesis<br>Marks an RA synovial tissue enriched peripheral T helper cell;<br>Marker of effector helper memory T cell |
| Kpna4 | 3.1673 | 4.00E-43 | Involved in NF-kB nuclear translocation |
| Mdm2 | 2.4483 | 9.82E-41 | Pro-arthritis in RA and CIA |
| Stat3 | 1.7648 | 2.93E-36 | Important in RA pathogenesis;<br>Possible treatment target;<br>Interaction with HIF1a |
| Pgf | 13.4457 | 5.96E-34 | Pro-RA;<br>Pro-angiogenesis |
| Atf4 | 1.6959 | 9.85E-34 | Pro-osteoclastogenic |
| Irak2 | 2.3285 | 3.83E-30 | Genetic risk factor in RA;<br>Pro-inflammatory |
| Hmga1 | 1.7851 | 5.00E-30 | Pro-angiogenesis;<br>Pro-inflammatory |
| Ptpn2 | 1.8440 | 7.19E-29 | RA- and JIA associated gene |
| Cd83 | 3.1311 | 5.70E-28 | RA-associated |
| Adam8 | 2.5072 | 2.71E-27 | Pro-osteoclastic;<br>Pro-inflammatory;<br>Pro-angiogenic |
| Jak1 | 1.8108 | 2.79E-25 | Therapeutically targeted RA signaling pathway |
| Sgk1 | 2.7032 | 4.77E-23 | Pro-Th17 |
| Il7r | 2.1721 | 1.84E-21 | Pro-inflammatory B cell |
| Relb | 1.8324 | 4.39E-18 | NF-kB factor associated with RA |
| Egr1 | 3.0682 | 4.51E-18 | Pro-angiogenic |
| Runx1 | 1.6525 | 6.03E-18 | Th17-polarizing |
| Bhlhe40 | 2.1067 | 1.91E-16 | Pro-angiogenic;<br>Pro-inflammatory |
| Vegfc | 2.3484 | 2.44E-15 | RA and angiogenesis associated |
| Traf1 | 2.1511 | 5.46E-15 | RA risk locus |
| Il1a | 5.2812 | 8.42E-15 | Pro-osteoclastogenic;<br>Therapeutically targetable in RA |
| Nxn | 2.9583 | 3.77E-13 | Pro-inflammatory |
| Vegfa | 1.9826 | 7.44E-13 | Angiogenesis, RA;<br>Angiogenesis and osteoclast-dependent bone remodeling;<br>Pro-osteoclastogenic |
| Ccr1 | 2.2640 | 2.58E-12 | Overexpressed in CIA joint;<br>Overexpressed in RA; |

| | | | |
|---|---|---|---|
| | | | Potentially targetable in RA |
| Tnfaip3 | 2.2224 | 3.27E-12 | RA risk locus |
| Jak3 | 1.8332 | 4.58E-12 | A pathogenic enzyme in RA;<br>Therapeutically targeted by tofacitinib |
| Src | 1.5709 | 1.71E-11 | Mediates IL-6 production in synovial fibroblasts;<br>Pro-osteoclast; Anti-osteoblast |
| Klf6 | 1.7604 | 2.37E-11 | Pro-M1 polarization |
| Cxcr4 | 2.4499 | 1.19E-09 | Involved in RA and angiogenesis;<br>Therapeutically targetable |
| Hk2 | 1.6686 | 1.25E-09 | Specifically expressed in RA and a proposed therapeutic target;<br>Pro-Th17; Pro-arthritis; Pro-angiogenic |
| Hbegf | 3.7274 | 3.54E-09 | IL-17-activated;<br>Pro-angiogenic |
| Smad6 | 2.1062 | 1.16E-08 | Pro-angiogenic |
| Gata3 | 1.8774 | 2.52E-08 | Pro-angiogenic |
| Dcstamp | 1.5013 | 5.67E-08 | Pro-osteoclastogenic |
| Vim | 1.7084 | 4.29E-07 | Considered an autoantigen in RA |
| Tnfaip6 | 4.5943 | 4.24E-06 | Pro-angiogenic |
| Mmp14 | 3.2325 | 4.38E-06 | Expressed in RA synovium;<br>Plays a role in joint destruction;<br>Therapeutically targetable |
| Olr1 | 4.0542 | 1.29E-05 | Pro-angiogenic |
| Tgfb3 | 2.6955 | 3.02E-05 | Pro-angiogenic |
| Mmp12 | 2.7932 | 4.31E-05 | Pathogenic role in inflammatory arthritis |
| Mmp13 | 2.9020 | 3.34E-04 | Overexpressed;<br>Plays a role in joint destruction |
| Cxcr1 | 3.0881 | 4.71E-04 | Transduces IL-8 pro-osteoclastogenic signal;<br>Pro-angiogenic |
| Egr2 | 1.9622 | 5.10E-04 | Risk factor for RA;<br>Upregulated by TNF |
| Sphk1 | 2.4153 | 3.65E-03 | Over-expressed in RA;<br>Pro-FLS activation;<br>Pro-angiogenic;<br>Proposed therapeutic target;<br>Downstream enzyme in the TNFa pathway |
| Areg | 2.0454 | 7.52E-03 | Over-expressed in RA synovium; Up-regulated by IL-1b;<br>Stimulates SFL in RA |
| Nanos1 | 1.5502 | 1.03E-02 | Increased expression in CIA;<br>Pro-angiogenic;<br>Pro-M1 polarization;<br>Pro-inflammatory |

| **Downregulated** | | | |
|---|---|---|---|
| | **Fold Change** | **Padj** | **Relevant roles/functions** |
| Abcb1b | 1.7458 | 3.41E-38 | Decreased levels in experimental arthritis rats |
| Akap1 | 1.6996 | 1.66E-50 | Ant-inflammatory;<br>Anti-hypoxia-induced stress |
| Anxa3 | 1.7587 | 1.61E-33 | RA susceptibility locus |
| Arhgap30 | 1.5626 | 3.90E-56 | Inhibitor of Wnt/β-catenin pathway |

| | | | |
|---|---|---|---|
| Atp2a3 | 2.1019 | 3.98E-41 | Its inhibition leads to NF-kB p65 translocation and improved survival of CD4 T cells |
| Bag2 | 1.7373 | 1.42E-26 | Repressed by NF-kB |
| Casp9 | 1.7946 | 2.62E-27 | Pro-apoptotic; anti-inflammatory |
| Cd81 | 1.6682 | 2.86E-24 | Pro-Th2 differentiation;<br>Anti-angiogenic |
| Chd4 | 1.5142 | 4.45E-29 | Mi-2. A co-repressor of T, B and plasma cell development |
| Ctdsp1 | 1.7170 | 1.81E-36 | Anti-angiogenic |
| Cyp51 | 2.2615 | 1.50E-41 | Anti-inflammatory - KO leads to inflammation |
| Dock2 | 1.5969 | 3.02E-64 | Immune regulatory |
| Fads1 | 1.6000 | 4.19E-77 | RA risk association;<br>Supporting M2 differentiation |
| Fkbp11 | 3.1032 | 1.17E-19 | Anti-ER stress and inflammation;<br>Pro-osteogenic |
| Glo1 | 1.7115 | 1.24E-29 | Part of HLA extended haplotype;<br>Anti-oxidant: downstream of Nrf2 |
| Hsd17b4 | 1.8641 | 1.31E-75 | Anti-inflammatory |
| Ikbip | 2.0823 | 6.82E-27 | Pro-apoptotic |
| Klf2 | 1.8713 | 2.40E-10 | Pro-M2;<br>Anti-angiogenic;<br>Anti-inflammatory |
| Lpcat3 | 1.9358 | 1.34E-36 | Pro-M2 polarization;<br>Anti-inflammatory |
| Lyl1 | 1.9867 | 2.95E-26 | Anti-angiogenic |
| Mapk14 | 1.6445 | 8.34E-30 | Anti-angiogenesis |
| Parp1 | 1.5292 | 9.48E-45 | Anti-osteoclastogenic |
| Patz1 | 1.8747 | 1.11E-27 | Anti-angiogenesis;<br>Inhibiting NF-kB pathway |
| Pi16 | 2.7190 | 2.22E-10 | Inhibits MMP2;<br>Treg subset marker |
| Pon2 | 1.9140 | 1.45E-35 | Pro-M2;<br>Anti-inflammatory;<br>Anti-oxidant;<br>Anti-atherogenic |
| Qdpr | 1.6821 | 4.49E-27 | Anti-oxidative damage |
| Scarb2 | 1.7086 | 2.80E-31 | CD36;<br>Anti-inflammatory |
| Slc13a3 | 2.4445 | 2.05E-09 | Anti-oxidative stress |
| Sub1 | 1.9104 | 4.56E-32 | Anti-oxidative stress |
| Txndc5 | 2.1280 | 1.26E-33 | Anti-oxidative stress |
| Ywhab | 1.6189 | 5.95E-61 | Anti-angiogenic; Associated with resolution of inflammation and M2 differentiation |

##### S1B. Notable unique DEGs modulated by 65-79*PE under M1 polarizing conditions

| **Upregulated** | | | |
|---|---|---|---|
| | **Fold Change** | **Padj** | **Relevant roles/functions** |
| Acox2 | 2.3960 | 1.46E-03 | Anti-oxidative stress |
| Alkbh2 | 1.5265 | 1.56E-12 | Post-inflammatory DNA damage-repairing enzyme |
| Bglap | 3.0692 | 7.84E-05 | Pro-osteoblast |
| Cdsn | 2.9308 | 2.04E-06 | Anti-IL-1b, anti-inflammatory |

| | | | |
|---|---|---|---|
| Def6 | 1.6748 | 2.58E-21 | Anti-osteoclastogenic;<br>Limits proliferation of TFH cells in mice via alteration of mTORC1 signaling |
| Dock3 | 2.6558 | 6.72E-07 | Anti-oxidative stress |
| Fbxl12 | 1.7321 | 1.01E-10 | pro-osteoblast |
| Filip1l | 1.8170 | 4.62E-15 | Anti-angiogenesis |
| Fyn | 1.6248 | 5.26E-13 | Anti-angiogenesis;<br>Anti-ER stress |
| Gpr65 | 1.7215 | 1.90E-14 | Anti-osteoclastogenic |
| Havcr2 | 1.7619 | 3.13E-21 | Pro-immune tolerance;<br>Anti-RA, inversely correlated with disease activity |
| Htatip2 | 1.6956 | 2.52E-12 | Anti-angiogenesis |
| Il17rc | 1.7079 | 6.22E-16 | Anti-IL-17<br>(in a soluble IL-17RC form) |
| Inpp4b | 2.5990 | 1.09E-03 | Anti-angiogenic;<br>Anti-osteoclast |
| Klhl22 | 1.6139 | 1.20E-30 | Activates mTORc |
| Mfsd2a | 2.1315 | 6.36E-07 | Anti-inflammatory;<br>Anti-angiogenic |
| Mt2 | 1.8269 | 1.44E-23 | Anti-oxidant;<br>Anti-inflammatory;<br>Inhibits NF-kB activation |
| Nrf1 | 1.5208 | 1.11E-10 | Anti-inflammatory;<br>Anti-m1 polarization |
| Ogg1 | 1.6344 | 1.51E-13 | Anti-inflammatory |
| Osgin1 | 1.6636 | 2.40E-11 | Cyto-protective, Nrf2-transcriptional target |
| Pdcd2 | 1.7995 | 4.16E-24 | Inhibits TNFa |
| Pycr1 | 1.8976 | 7.72E-11 | Anti-oxidative stress |
| Stc2 | 1.9779 | 1.70E-11 | Anti-ER and oxidative stress |
| Stk10 | 1.6032 | 1.14E-10 | Anti-inflammatory;<br>Anti-lymphocyte activation and inflammation |
| Themis2 | 1.8008 | 3.61E-22 | Inhibits macrophage activation |
| Tlr1 | 1.5914 | 6.78E-17 | Pro-M2 |
| Traf3ip3 | 1.8829 | 1.29E-25 | Regulator of Treg;<br>Interacts with mTORC1 |

| **Downregulated** | | | |
|---|---|---|---|
| | **Fold Change** | **Padj** | **Relevant roles/functions** |
| Atp6v0d2 | -2.7543 | 1.49E-48 | Pro-osteoclastogenic |
| Oas2 | -2.6083 | 2.59E-40 | RA signature gene regulating immune response;<br>Innate, type 1 IFN-induced |
| Dnmt3a | -2.0215 | 7.63E-35 | Pro-osteoclastogenic;<br>Over-expressed in RA synovium;<br>Expression in macrophages directly correlates with pro-inflammatory and inversely with anti-inflammatory state |
| St3gal1 | -2.7146 | 5.71E-33 | Pro-angiogenic |
| Syk | -2.1744 | 3.26E-31 | Pro-arthritogenic in RA model;<br>Therapeutic target in RA;<br>Pro-osteoclastogenic |
| Colec12 | -2.5837 | 1.16E-29 | Pro-inflammatory |
| Glul | -2.0115 | 3.84E-27 | Pro-angiogenic |
| Epn2 | -1.9301 | 6.75E-27 | Pro-angiogenic |

| | | | |
|---|---|---|---|
| Ckb | -2.1008 | 2.91E-26 | Pro-osteoclastogenic |
| Sh3kbp1 | -2.0065 | 6.88E-25 | NF-kB activation;<br>B and T cell immune response |
| Ddx58 | -1.8876 | 1.69E-23 | Innate immune system;<br>Activator of NF-kB and IRF-3;<br>Abundant in RA synovial fibroblasts |
| Jun | -2.8318 | 4.77E-23 | Increased expression in activated synovial cells and in patients;<br>Pro-arthritis in mice |
| Gtf2i | -1.5113 | 3.26E-22 | RA-associated locus in Asians;<br>Autoimmunity-associated locus |
| Tnfrsf9 | -4.5323 | 4.43E-22 | Released by activated lymphocytes and is abundant in RA;<br>Therapeutically targetable in arthritic mice; RA severity locus in African Americans |
| Tspan5 | -1.6426 | 1.19E-21 | Pro-osteoclastic |
| Lrp1 | -2.2938 | 1.43E-21 | Co-receptor of SE ligand 65-79*0401 |
| Bhlhe41 | -1.9412 | 1.52E-20 | Abundant in RA synovium, increases IL1b |
| F10 | -5.0286 | 4.06E-20 | Factor X. Potential key role in RA progression |
| F7 | -4.0622 | 6.75E-09 | Factor VII. Plays a role in the pathogenesis of chronic destructive arthritis in RA; NF-Kb activator and IL-8 inducer; Inducer of VEGF in fibroblasts; Pro-angiogenic |
| Arhgap24 | -3.0636 | 1.17E-07 | Rac1. Plays a role in RA synoviocyte activation;<br>Pro-arthritogenic in mice;<br>Pro-angiogenic |
| Sulf2 | -3.0302 | 4.16E-07 | Expressed in RA synoviocytes and involved in their activation state; Proangiogenic |
| Acp5 | -2.4707 | 6.03E-07 | Tartrate resistant acid phosphatase - a key plyer in osteoclast-mediated bone erosion |
| Scg2 | -3.2210 | 8.12E-07 | Pro-angiogenic |
| Fabp4 | -2.7021 | 2.42E-06 | Increased levels in RA;<br>Pro-angiogenic |
| Adamts7 | -2.5569 | 4.48E-04 | Over-expressed in RA joint tissues and degrades cartilage;<br>Arthritogenic in mice |
| Src | -1.5104 | 2.94E-09 | Pro-osteoclast |

#### S1C. Notable unique DEGs modulated by 65-79*SE under M2 polarizing conditions

| **Upregulated** | | | |
|---|---|---|---|
| | **Fold Change** | **Padj** | **Relevant roles/functions** |
| Slc31a2 | 1.7121 | 1.09E-10 | Pro-oxidative |
| Adamtsl5 | 1.5423 | 1.07E-09 | Inducer of IL-17 |
| Ryr1 | 1.5904 | 3.06E-07 | Pro-oxidative;<br>NF-kB inducible |
| Layn | 1.6915 | 8.93E-07 | Enhances inflammation and cartilage destruction and secretion of IL-8 |
| Mmp9 | 2.0024 | 1.43E-06 | RA;<br>Pro-osteoclast;<br>Pro-angiogenesis |
| Trpc4 | 1.6466 | 2.31E-06 | Pro-angiogenic |
| Lat | 1.9123 | 1.56E-05 | RA |

| | | | |
|---|---|---|---|
| Atf3 | 1.5921 | 2.14E-05 | Pro-angiogenic |
| Il11ra1 | 1.6034 | 5.37E-05 | Pro-osteoclast and bone remodeling |
| Ahnak2 | 1.5733 | 6.13E-05 | Associated with RA risk |
| Lcn2 | 1.6957 | 7.38E-05 | Associated with RA |
| Fas | 1.6118 | 3.50E-04 | Associated with RA |
| Anpep | 1.6007 | 4.24E-04 | CD13 |
| Pgf | 1.6470 | 5.30E-04 | Pro-RA;<br>Pro-angiogenesis |
| Arg2 | 1.5223 | 1.00E-03 | M1 macrophage marker;<br>Correlates with RA disease activity;<br>Pro-oxidative |
| Src | 1.5127 | 1.45E-03 | Pro-osteoclast |
| Ccl7 | 1.6488 | 1.78E-03 | Activates monocyte migration in RA |
| Acod1 | 1.5991 | 2.33E-03 | Pro-inflammatory;<br>M1 macrophage marker |
| Atp6v0d2 | 1.6287 | 2.43E-03 | Pro-osteoclastogenic |
| Cd74 | 1.5001 | 2.64E-03 | MIF in RA |
| Cd84 | 1.5074 | 5.95E-03 | RA risk factor and disease marker |

| **Downregulated** | | | |
|---|---|---|---|
| | **Fold Change** | **Padj** | **Relevant roles/functions** |
| Rnase4 | 1.7690 | 6.96E-10 | Regulator of innate immune response |
| Col18a1 | 1.6365 | 2.18E-08 | Anti-angiogenesis;<br>Proposed treatment for RA |
| Stk17b | 1.5931 | 2.27E-06 | Anti-memory T cell development; Anti-T cell activation |
| Zfp608 | 1.8395 | 2.59E-06 | Anti-thymocytes - prevents RAG1 and RAG2 |
| Cd276 | 1.5893 | 1.60E-05 | M2-polarizing;<br>Inhibitor of T cell response |
| Txnip | 1.5641 | 4.18E-04 | Inhibitor of HIF1a |
| Slc8a1 | 1.6686 | 4.27E-04 | Inhibitor of osteoclast-mediated bone resorption |
| Nt5e | 1.5624 | 1.74E-03 | Protective against CIA;<br>promotes M2;<br>Marker of Treg and mediates immune suppression |
| Cx3cr1 | 1.6004 | 2.95E-03 | M2 marker in mice;<br>Required for Treg proliferation and IL-10 production |

#### S1D. Notable unique DEGs modulated by 65-79*PE under M2 polarizing conditions

| **Upregulated** | | | |
|---|---|---|---|
| | **Fold Change** | **Padj** | **Relevant roles/functions** |
| Glrx | 1.7104 | 2.88E-19 | Anti-redox; anti-angiogenesis |
| Anxa6 | 1.5611 | 9.59E-13 | NF-kB modulator |
| Car5b | 1.5519 | 3.09E-12 | Attenuated macrophages by PCP |
| Anxa1 | 1.5247 | 5.37E-11 | Anti-inflammatory in arthritis |
| Rgs2 | 1.8427 | 1.38E-09 | Anti-inflammatory |
| Col7a1 | 1.8549 | 3.06E-08 | Anti angiogenic |
| Ctss | 1.5120 | 1.59E-07 | JNK1/2 pathway modulator:<br>pro-osteoblast, anti-bone remodeling |
| Rras | 1.5516 | 1.14E-05 | Activates PI3K-Akt; Anti-angiogenic |
| Fcna | 1.5001 | 1.34E-05 | RA-associated locus polymorphism:<br>Negative correlation with SLE disease activity |

| | | | |
|---|---|---|---|
| Lamb2 | 1.7026 | 1.62E-05 | Pro-angiogenic |
| Spn | 1.5961 | 2.17E-05 | Increases IL4R and IL4;<br>Anti-monocyte and anti-T cell; adhesion; Inhibits NF-kB activation;<br>T cell-constraining activity |
| Pdzd2 | 1.8577 | 6.25E-05 | Response to ant-TNF treatment locus |
| Glipr2 | 1.7753 | 1.51E-04 | Modulates cytokine production; Negative regulator of autophagy |
| Tnfrsf17 | 1.7698 | 3.57E-04 | Immunosuppressing |

| **Downregulated** | | | |
|---|---|---|---|
| | **Fold Change** | **Padj** | **Relevant roles/functions** |
| Arid3a | 1.6710 | 3.30E-12 | Marker of CD19+ B lymphocytes |
| Cd80 | 1.6373 | 1.19E-05 | Therapeutic target for RA |
| Cxcl10 | 1.7267 | 8.51E-05 | Marker for RA disease and activity |
| Cxcl2 | 1.7056 | 1.55E-04 | RA associated;<br>NF-kB-mediated osteoclastogenesis |
| Cxcr3 | 1.5758 | 1.84E-03 | CXCL10 receptor, marker for RA disease and activity |
| Fam102a | 1.8050 | 2.58E-12 | Pro-osteoclastogenic |
| Fcrl1 | 1.5430 | 3.07E-11 | Association with RA |
| Ifi44l | 2.2361 | 1.04E-10 | Interferon-inducible gene associated with RA |
| Il21r | 1.7233 | 1.41E-04 | RA pathogenesis |
| Myo1d | 1.7571 | 3.22E-09 | Pro-osteoclastogenic |
| Nfkbiz | 1.5678 | 1.80E-04 | Th17-differentiating transcription factor |
| Osgin1 | 1.7734 | 1.72E-08 | Oxidative stress inducible autophagy |
| Prkca | 1.6536 | 6.72E-10 | Pro-angiogenic |
| Rsad2 | 1.7053 | 1.51E-04 | RA-associated and type I interferon response gene |
| Slc40a1 | 2.1273 | 1.28E-09 | Pro-inflammatory |
| Tnf | 1.7878 | 3.91E-07 | Therapeutic target for RA;<br>Activation of osteoclasts |
| Traf3ip3 | 1.6060 | 1.75E-09 | Regulator of T and B cell development |

### Table S2: Notable upstream regulators with autoimmune disease relevance

#### S2A. Upstream regulators for 65-79*SE under M1 polarizing conditions

| Upstream Regulator | Relevant functions | Predicted | z-score | p-value |
|---|---|---|---|---|
| **STAT3** | Pro-Th cells in RA; Pro-arthritis in mice; Pro-synoviocyte survival in RA | **Activated** | 2.03 | 3.51E-35 |
| **JUN** | Overexpressed in RA; Pro-arthritis in mice | **Activated** | 2.874 | 1.45E-21 |
| **EGR1** | Pro-osteoclastogenic; NF-κB activator | **Activated** | 3.129 | 4.29E-19 |
| **SP1** | IL-1a activator; Induction of CCL2 expression | **Activated** | 2.156 | 3.14E-17 |
| **EP300** | Induction of CCL2 and IL-6 expression: Mediates IL-1β pro-inflammatory effect on RA synoviocytes | **Activated** | 2.05 | 3.61E-17 |
| **RELA** | Pathogenic factor in RA; Pro-arthritis in mice; Pro-osteoclastogenic | **Activated** | 4.198 | 3.06E-15 |
| **CEBPB** | Pro-inflammatory | **Activated** | 3.455 | 3.27E-15 |
| **NUPR1** | Pro-IL-1β-mediated expression of MMP13; Activates the NF-κB pathway | **Activated** | 2.494 | 5.08E-15 |
| **CTNNB1** | Activates RA synoviocytes; Pro-osteoclastogenic | **Activated** | 3.661 | 7.14E-12 |
| **SMAD3** | Pro-osteoclastogenic | **Activated** | 3.119 | 1.71E-11 |
| **REL** | Pathogenic factor in RA; RA susceptibility locus | **Activated** | 3.323 | 7.71E-11 |
| **CEBPA** | Inhibits M2 macrophage differentiation | **Activated** | 2.812 | 9.03E-11 |
| **HIF1A** | Pro-inflammatory in RA; Pro-angiogenic in RA; Activates PAD2-mediated protein citrullination in RA | **Activated** | 2.42 | 9.62E-11 |
| **KLF2** | Pro-M2 macrophage differentiation; Anti-arthritis in mice; Anti-angiogenic | **Inhibited** | -2.166 | 1.77E-10 |
| **FOXP3** | Pro-Treg; RA-protective | **Inhibited** | -2.96 | 1.78E-09 |
| **XBP1** | Anti-inflammatory; NF-κB inhibitor | **Inhibited** | -2.048 | 5.7E-10 |

#### S2B. Upstream regulators for 65-79*PE under M1 polarizing conditions

| Upstream Regulator | Relevant functions | Predicted | z-score | p-value |
|---|---|---|---|---|
| **SREBF1** | Pro-inflammatory; Over-expressed in RA synoviocytes; Pro-angiogenic | **Inhibited** | -2.034 | 2.61E-10 |
| **KEAP1** | Pro-inflammatory in basal conditions by promoting degradation of Nrf2 | **Inhibited** | -2.191 | 5.1E-07 |
| **MITF** | Pro-osteoclastogenic | **Inhibited** | -2.814 | 0.000165 |
| **SOX11** | Pro-angiogenic | **Inhibited** | -2.292 | 0.000202 |
| **E2F1** | Pro-osteoclastogenic; Pro-angiogenic | **Inhibited** | -2.353 | 0.000401 |
| **NFKBIZ** | Pro-Treg; Anti-inflammatory: Enhances IL10 expression | **Activated** | 2.122 | 0.00178 |

| | | | | |
|---|---|---|---|---|
| MTA1 | Modulates cytokine networks in RA; Anti-inflammatory | **Activated** | 2.378 | 0.00938 |
| TARDBP | Inhibits NF-κB activity; Anti-inflammatory | **Activated** | 2.236 | 0.0301 |
| XBP1 | Anti-inflammatory; NF-κB inhibitor | **Activated** | 2.401 | 0.0492 |

#### S2C. Upstream regulators for 65-79*SE under M2 polarizing conditions

| Upstream Regulator | Relevant functions | Predicted | z-score | p-value |
|---|---|---|---|---|
| EGR1 | Over-expressed in RA synoviocytes; Pro-inflammatory in RA synoviocytes; Pro-angiogenic | **Activated** | 2.229 | 2.36E-06 |
| EPAS1 | Pro-angiogenic | **Activated** | 2.382 | 0.00396 |
| GLI1 | Pro-inflammatory cytokines in RA; Pro-RA synoviocyte proliferation | **Activated** | 2.371 | 0.0148 |
| IRF3 | Activates RA synoviocytes; Activates antibody production by B cells; Pro-M1; Pro-angiogenic | **Activated** | 2.414 | 0.000152 |
| IRF7 | Pro-M1; Pro-arthritogenic in CIA mice | **Activated** | 2.236 | 0.0257 |
| TCF3 | Inhibits the pro-RA β-catenin/Wnt pathway | **Inhibited** | -2 | 0.00555 |
| RUNX1 | Anti-osteoclast; Anti-angiogenesis | **Inhibited** | -2 | 0.00947 |

#### S2D. Upstream regulators for 65-79*PE under M2 polarizing conditions

| Upstream Regulator | Relevant functions | Predicted | z-score | p-value |
|---|---|---|---|---|
| STAT6 | Pro-M2; Anti-arthritis in mice | **Activated** | 2.075 | 1.84E-11 |
| KLF2 | Pro-M2; Anti-arthritis in mice; Anti-angiogenic | **Activated** | 2.401 | 0.0000233 |
| NFATC2 | Pro-osteoclastogenic; Pro-angiogenic | **Inhibited** | -2.091 | 1.82E-11 |
| IRF5 | Pro-inflammatory; RA-associated; RA risk locus; Associated with erosive RA; Pro-arthritis in mice | **Inhibited** | -2.362 | 0.0000356 |
| EZH2 | Over-expressed in RA synovium; Pro-osteoclastogenic; Pro-angiogenic | **Inhibited** | -2.601 | 0.000131 |
| GATA6 | Pro-angiogenic | **Inhibited** | -2.013 | 0.000144 |
| KLF6 | Pro-M1 and pro-inflammatory; Pro-angiogenic | **Inhibited** | -2.131 | 0.00184 |
| IRF3 | Activates RA synoviocytes; Activates antibody production by B cells; Pro-M1; Pro-angiogenic | **Inhibited** | -2.624 | 1.84E-07 |
| IRF7 | Pro-M1; Pro-arthritogenic in CIA mice | **Inhibited** | -2.115 | 2.74E-09 |

Purple colored regulators indicate pro-inflammatory or pro-RA regulators; green colored regulators indicate anti-inflammatory or anti-RA regulators. Yellow highlight identifies regulators with opposite activation status in 65-79*SE vs 65-79*PE

### Table S3: List of reagents used in this study.

| Reagent or Resource | Source | Category numbers |
|---|---|---|
| **Antibodies** | | |
| Anti-mouse Akt | Cell Signaling Technologies (Danvers, MA) | Cat# 9272 |
| Anti-mouse Phospho-Akt | Cell Signaling Technologies (Danvers, MA) | Cat# 9271 |
| Anti-mouse Stat6 | Cell Signaling Technologies (Danvers, MA) | Cat# 9362 |
| Anti-mouse Phospho-Stat6 | Cell Signaling Technologies (Danvers, MA) | Cat#56554 |
| Anti-mouse SHIP1 | Cell Signaling Technologies (Danvers, MA) | Cat# 2728 |
| Anti-mouse Phospho-SHIP1 | Cell Signaling Technologies (Danvers, MA) | Cat# 3941 |
| Anti-mouse PTEN | Cell Signaling Technologies (Danvers, MA) | Cat# 9559 |
| Anti-mouse Phospho-PTEN | Cell Signaling Technologies (Danvers, MA) | Cat# 9549 |
| Anti-mouse beta-actin | Invitrogen (Waltham, MA) | Cat# BA3R |
| Anti-rabbit IgG HRP-linked | Cell Signaling Technologies (Danvers, MA) | Cat# 7074 |
| Anti-mouse IgG HRP-linked | GE healthcare Lifesciences (Chicago, IL) | Cat# NA931 |
| **Biological Samples** | | |
| Fetal Bovine Serum | Corning (Tewksbury, MA) | Cat# 35-015-CV |
| Antibiotics (Pen Strep) | Gibco (Waltham, MA) | Cat# 15140-122 |
| **Media** | | |
| DMEM | Gibco (Waltham, MA) | Cat# 11965-092 |
| DMEM | Gibco (Waltham, MA) | Cat# 11885-084 |
| DMEM | Sigma (St. Louis, MO) | Cat# D5921 |
| MEM Alpha | Gibco (Waltham, MA) | Cat# 12561-056 |
| RPMI 1640 | Gibco (Waltham, MA) | Cat# 11875-093 |
| **Cells** | | |
| RAW 264.7 | ATCC (Manassas, VA) | Cat# TIB-71 |
| THP-1 cells | ATCC (Manassas, VA) | Cat# TIB-202 |
| **Mice** | | |
| HLA-DRB1*0401 | Chela David | (*35*) |
| HLA-DRB1*0402 | Chela David | (*26*) |
| **Chemicals, Reagents, peptides, and Recombinant Proteins** | | |
| Recombinant Murine Il-4 | R&D (Minneapolis, MN) | Cat# 404-ML-010 |
| Recombinant Murine Il-13 | R&D (Minneapolis, MN) | Cat# 413-ML-010 |
| Recombinant Murine IFNγ | Peprotech (Rocky Hill, NJ) | Cat# 315-05 |
| Recombinant Murine MCSF | Peprotech (Rocky Hill, NJ) | Cat# 315-02 |
| Recombinant Human IFNγ | Peprotech (Rocky Hill, NJ) | Cat# 300-02 |
| LPS | Sigma (St. Louis, MO) | Cat# L2630 |
| 65-79*SE (65-79*0401) | Bioworld (Dublin, OH) | (*16*) |
| 65-79*PE (65-79*0402) | Bioworld (Dublin, OH) | (*16*) |
| PBS (Phosphate buffered saline) | Gibco (Waltham, MA) | Cat# 10010-023 |
| Bovine Serum Albumin | Sigma (St. Louis, MO) | Cat# A7906 |
| Sodium Pyruvate | Gibco (Waltham, MA) | Cat# 11360-070 |
| DAF-2 DA | Sigma-Millipore (St. Louis, MO) | Cat# 251505 |

| | | |
|---|---|---|
| Novex™ 4-20% Tris-Glycine gels | Invitrogen (Waltham, MA) | Cat# XP04200BOX |
| RIPA Buffer | Sigma (St. Louis, MO) | Cat# R0278-50ml |
| cOmplete mini EDTA-free | Roche (Indianapolis, IN) | Cat# 11836170001 |
| phosSTOP | Roche (Indianapolis, IN) | Cat# 04906845001 |
| Trizol | Thermo Fisher (Waltham, MA) | Cat# 155596018 |
| SuperSignal$^{TM}$ West Pico Plus ECL substrate | Thermo Scientific (Waltham, MA) | Cat# 34577 |
| Ly294002 | Sigma (St. Louis, MO) | Cat# L9908 |
| Wedelolactone | Sigma (St. Louis, MO) | Cat# W4016 |
| PF-4708671 | Selleckchem (Houston, TX) | Cat# S2163 |
| Phorbol 12-myristate 13-acetate | Sigma (St. Louis, MO) | Cat# P8139 |
| See Supplementary materials for the Primer sequences used for qPCR | Integrated DNA Technology (Coralville, IA) | |
| **Critical commercial kits and assays** | | |
| Mouse TNF-alpha DuoSet ELISA | R&D (Minneapolis, MN) | Cat# Dy-410-05 |
| Mouse Il-12p70 DuoSet ELISA | R&D (Minneapolis, MN) | Cat# Dy-419-05 |
| Mouse Il-6 DuoSet ELISA | R&D (Minneapolis, MN) | Cat# Dy-406-05 |
| Arginase Activity Colorimetric Assay Kit | Biovision Inc. (Milpas, CA) | Cat# K755 |
| Direct-zol$^{TM}$ RNA MiniPrep | Zymo Research (Irvine, CA) | Cat# R2052 |
| RNeasy Plus Mini kit | Qiagen (Germantown, MD) | Cat# 74134 |
| TruSeq RNA Sample Prep Kit | Illumina (San Diego, CA) | Cat# RS-122-2001 |
| Fast SYBR$^{TM}$ Green Master Mix | Thermo Fisher (Waltham, MA) | |
| High Capacity cDNA Reverse transcription Kit | Thermo Fisher (Waltham, MA) | |
| TURBO DNA-free™ Kit | Invitrogen (Waltham, MA) | Cat# AM1907 |
| Quantibody® Mouse Cytokine Array 1 | Biolegend (San Diego, CA) | Cat# QAM-CYT-1 |
| *DC* protein assay Kit | BioRad (Hercules, CA) | Cat# 5000112 |
| **Software and Algorithms** | | |
| Prism V8.1.0 | GraphPad Software, Inc. | https://www.graphpad.com/scientificsoftware/prism/ |
| ImageJ | | https://imagej.nih.gov/ij/ |
| Rsubread1.5.0p3 | (*57*) | |
| DESeq2-1.16.1 | (*58*) | http://bioconductor.org/packages/release/bioc/html/DESeq2.html |
| DAVID 6.8 | https://david.ncifcrf.gov/ | (*55*) |
| Ingenuity Pathway Analysis | Qiagen (Germantown, MD) | https://www.qiagenbioinformatics.com/pr |

| | | |
|---|---|---|
| | | oducts/ingenuity-pathway-analysis |
| **Deposited data** | | |
| RNA-seq data | This paper | GEO |
| **Other** | | |
| Omega Lum C imaging system | Aplegen (San Francisco, CA) | |
| Synergy H1 Hybrid Multi-Mode Reader | Biotek (Winooski, VT) | https://www.biotek.com/products/detection-hybrid-technology-multi-mode-microplate-readers/synergy-h1-hybrid-multi-mode-reader/ |

#### Table S4: List of primer sequences used for qPCR analysis.

| Gene | Forward | Reverse |
|---|---|---|
| *Ccl2* | GAGGAAGGCCAGCCCAGCAC | TGGATGCTCCAGCCGGCAAC |
| *Il-12p40* | TCTTCTGCTTGGTTGGCTTT | CTCTGCGGGCATTTAACATT |
| *Cxcl10* | GGATGGCTGTCCTAGCTCTG | TGAGCTAGGGAGGACAAGGA |
| *Nos2* | CACCTTGGAGTTCACCCAGT | ACCACTCGTACTTGGGATGC |
| *Arg1* | GACCTGGCCTTTGTTGATGT | CAGCTCTTCATTGGCTTTCC |
| *Ym* | TAGTACTGGCCCACCAGGAA | AGACCTCAGTGGCTCCTTCA |
| *Il-10* | TGCACTACCAAAGCCACAAG | TAAGAGCAGGCAGCATAGCA |
| *Il-6* | CTTCACAAGTCCGGAGAGGA | TCCACGATTTCCCAGAGAAC |
| *Tnfa* | CTGGGACAGTGACCTGGACT | CTCCCTTTGCAGAACTCAGG |
| *Il-1b* | CAGGCAGGCAGTATCACTCA | TGTCCTCATCCTGGAAGGTC |
| *Il-12p35* | CATCGATGAGCTGATGCAGT | GAAGCAGGATGCAGAGCTTC |
| *Il-23p19* | CCAGCGGGACATATGAATCT | AGTCCTTGTGGGTCACAACC |
| *Mgl2* | ATCGCTTAGCCAATGTGCTT | TGGCCTCCAATTCTTGAAAC |
| *Ccl17* | AGTGGAGTGTTCCAGGGATG | TGGCCTTCTTCACATGTTTG |
| *Adgre1* | TGGAACACTGATGTGGAGGA | AGTTTGCCATCCGGTTACAG |
| *Hprt* | GCCCCAAAATGGTTAAGGTT | TTGCGCTCATCTTAGGCTTT |
